# Genome-wide patterns of homoeologous gene flow in allotetraploid coffee

**DOI:** 10.1101/2023.09.10.557041

**Authors:** Andre J. Ortiz, Joel Sharbrough

## Abstract

**Premise:** Allopolyploidy – a hybridization-induced whole-genome duplication event – has been a major driver of plant diversification. The extent to which chromosomes pair with their proper homolog vs. with their homoeolog in allopolyploids varies across taxa, and methods to detect homoeologous gene flow (HGF) are needed to understand how HGF has shaped polyploid lineages.

**Methods:** The ABBA-BABA test represents a classic method for detecting introgression between closely related species, but here we developed a modified use of the ABBA-BABA test to characterize the extent and direction of HGF in allotetraploid *Coffea arabica*.

**Results:** We found that HGF is abundant in the *C. arabica* genome, with both subgenomes serving as donors and recipients of variation. We also found that HGF is highly maternally biased in plastid-targeted – but not mitochondrial-targeted – genes, as would be expected if plastid-nuclear incompatibilities exist between the two parent species.

**Discussion:** Together our analyses provide a simple framework for detecting HGF and new evidence consistent with selection favoring overwriting of paternally derived alleles by maternally derived alleles to ameliorate plastid-nuclear incompatibilities. Natural selection therefore appears to shape the direction and intensity of HGF in allopolyploid coffee, indicating that cytoplasmic inheritance has long-term consequences for polyploid lineages.

## INTRODUCTION

Whole-genome duplication events (WGDs), in which the entire nuclear genome is doubled resulting in polyploid lineages, are widespread among plants, especially angiosperms (Jiao et al., 2011; Wendel, 2015; Ruprecht et al., 2017; One Thousand Plant Transcriptomes Initiative, 2019; Román-Palacios et al., 2021; Heslop-Harrison et al., 2023). WGDs are among the most profound mutational changes observed in nature particularly because they result in global genomic redundancy, which has consequences that range from the gene to the population (Fox et al., 2020). At the gene level, genomic redundancy contributes to relaxation of selective constraints (Otto and Whitton, 2000; Douglas et al., 2015; Zhang et al., 2020; Conover and Wendel, 2022), transcriptional re-programming (Schnable et al., 2011; Combes et al., 2013; Yoo et al., 2013; Akama et al., 2014; Hu et al., 2016; Yang et al., 2016; Edger et al., 2017; Ramírez-González et al., 2018; Oberprieler et al., 2019; Landis et al., 2020; Song et al., 2020), altered epigenetic regulation (Madlung et al., 2002; Salmon et al., 2005; Shcherban et al., 2008; Fulneček et al., 2009; Akagi et al., 2016; Chen et al., 2017; Song et al., 2017; Ding and Chen, 2018; Rao et al., 2023), transposable element expansion (Ågren et al., 2016; Baduel et al., 2019), altered rates of homologous, ectopic, and intergenomic recombination (Chalhoub et al., 2014; Guo et al., 2014; Jarvis et al., 2017; Chen et al., 2018; Bertioli et al., 2019; Mason and Wendel, 2020; Gonzalo et al., 2023), chromosomal structural changes (Chester et al., 2012; Edwards et al., 2017; Gordon et al., 2020; Cai et al., 2021; Orantes-Bonilla et al., 2022), among a host of other fundamental changes to genome biology (Otto, 2007; Leitch and Leitch, 2008; Doyle and Coate, 2019; Bomblies, 2020), all of which have the potential to directly impact organismal function and fitness.

Polyploids are generally categorized into one of two categories: allopolyploid or autopolyploid (Kihara and Ono, 1926; Soltis et al., 2014), depending upon whether their formation occurred via hybridization (Doyle et al., 2008) or via unreduced gametes (Mason and Pires, 2015). The mechanism of formation has important implications for post-WGD evolution, namely in the pattern of chromosome pairing during meiosis (Comai, 2005). Autopolyploids often exhibit multisomic inheritance in which homologs pair randomly, or even form tetravalents, during meiosis (Ramsey and Schemske, 2003). These unusual meiotic patterns are almost certainly part of the explanation for why so many polyploids also reproduce asexually (Otto and Whitton, 2000; Bomblies et al., 2016). By contrast, allopolyploids exhibit a range of different inheritance patterns ranging from completely disomic, in which homoeologs pair with their correct homolog during meiosis (*e.g*., cotton (Endrizzi, 1962)), to tetrasomic, in which homoeologs can pair with the correct homolog or with the homoeolog from the opposing subgenome (*e.g*., peanut (Leal-Bertioli et al., 2018)). The range of inheritance patterns exhibited by allopolyploid taxa can even be observed within the same genome (*e.g*., tobacco (Edwards et al., 2017)). Pairing and recombination between homoeologs from opposite subgenomes can even result in homoeologous exchange (cross-over) or homoeologous gene conversion (non-crossover), leading to gene flow across subgenomes (Mason and Wendel, 2020), and this process is expected to be especially prominent in relatively young allopolyploids (*e.g*., peanut, coffee). This homoeologous gene flow (HGF) therefore provides a mechanism for the production of novel genotypes that were absent from the constituent subgenomes at the time of polyploid formation. With the rise of long-read sequencing and high quality polyploid genomic resources, patterns of HGF and some of its phenotypic effects have been documented in a number of allopolyploid systems (Gaeta et al., 2007; Lashermes et al., 2016; Xiong et al., 2020; Zhang et al., 2020; Chu et al., 2021), although the contributions of variation derived from HGF to allopolyploid lineage success remains poorly understood (Deb et al., 2023).

As with inter-species introgression, HGF may provide allopolyploids access to adaptive variation that has already been tested by nature. Adaptive variation obtained by HGF may be particularly important for maintaining epistatic interactions in the face of genome mergers, as it provides a means to overcome Bateson-Dobzhansky-Muller incompatibilities (BDMIs) that can accompany genome merger (Sharbrough et al., 2017). That is, if HGF can facilitate replacement of incompatible components of epistatic interactions with compatible ones, allopolyploid lineages may be able to avoid some of the deleterious consequences of hybridization. One particularly useful example can be found in the interactions between nuclear-encoded and cytoplasmically encoded (*i.e*., mitochondria and chloroplast) genes and gene products, which are critical for carrying out the essential processes of respiration and photosynthesis (Rand et al., 2004; Sloan et al., 2018). In particular, multi-subunit enzyme complexes that are jointly encoded by the nuclear genome and the cytoplasmic genomes (*e.g*., Complexes I, III, IV, and V of the electron transport chain in mitochondria, and Rubisco, both Photosystems, and the CLP protease in chloroplasts, etc. (Forsythe et al., 2019) produce the lion’s share of the cell’s energy budget. Because the cytoplasmic genomes are usually inherited from only a single progenitor (Camus et al., 2022), while nuclear genomes are biparentally inherited from both progenitors, co-evolution between nuclear and cytoplasmic genes (*i.e*., cytonuclear co-evolution) in the maternal lineage can result in incompatibilities between the cytoplasmic genomes and the paternally derived half of the nuclear genome (Sharbrough et al., 2017). Maternally biased HGF provides an opportunity to ameliorate such cytonuclear incompatibilities in allopolyploids (just as co-introgression can act to maintain epistatic interactions in inter-species gene flow (Beck et al., 2015)), while other epistatic modules may benefit from HGF in either direction.

*Coffea arabica* (4x = 2n = 44) is an economically important allotetraploid crop contributing to ≥65% of global coffee consumption (∼170 million bags/year in total, https://www.ico.org/es/Market-Report-22-23-c.asp) (Campuzano-Duque et al., 2021), and is also an excellent model for studying HGF. In particular, *C. arabica* is the result of an hybridization event between *Coffea canephora* (2x = 2n = 22; paternal diploid progenitor) and *Coffea eugenioides* (2x = 2n = 22; maternal/cytoplasmic diploid progenitor) (Figure 1), with some debate over whether it evolved recently (*i.e.*, ∼10,000 – 50,000 years ago (Cros et al., 1998; Lashermes et al., 1999; Scalabrin et al., 2020) or more anciently (610,000 years ago – (Salojarvi et al., 2023)). Regardless, there has been sufficient time for HGF to occur, yet the maternal (E subgenome) and paternal (C subgenome) subgenomes remain distinguishable (mean *d_S_* ∼ 2.6%, mean *d_N_* ∼1.0% between diploids; (Sharbrough et al., 2022)).

**Figure 1.**
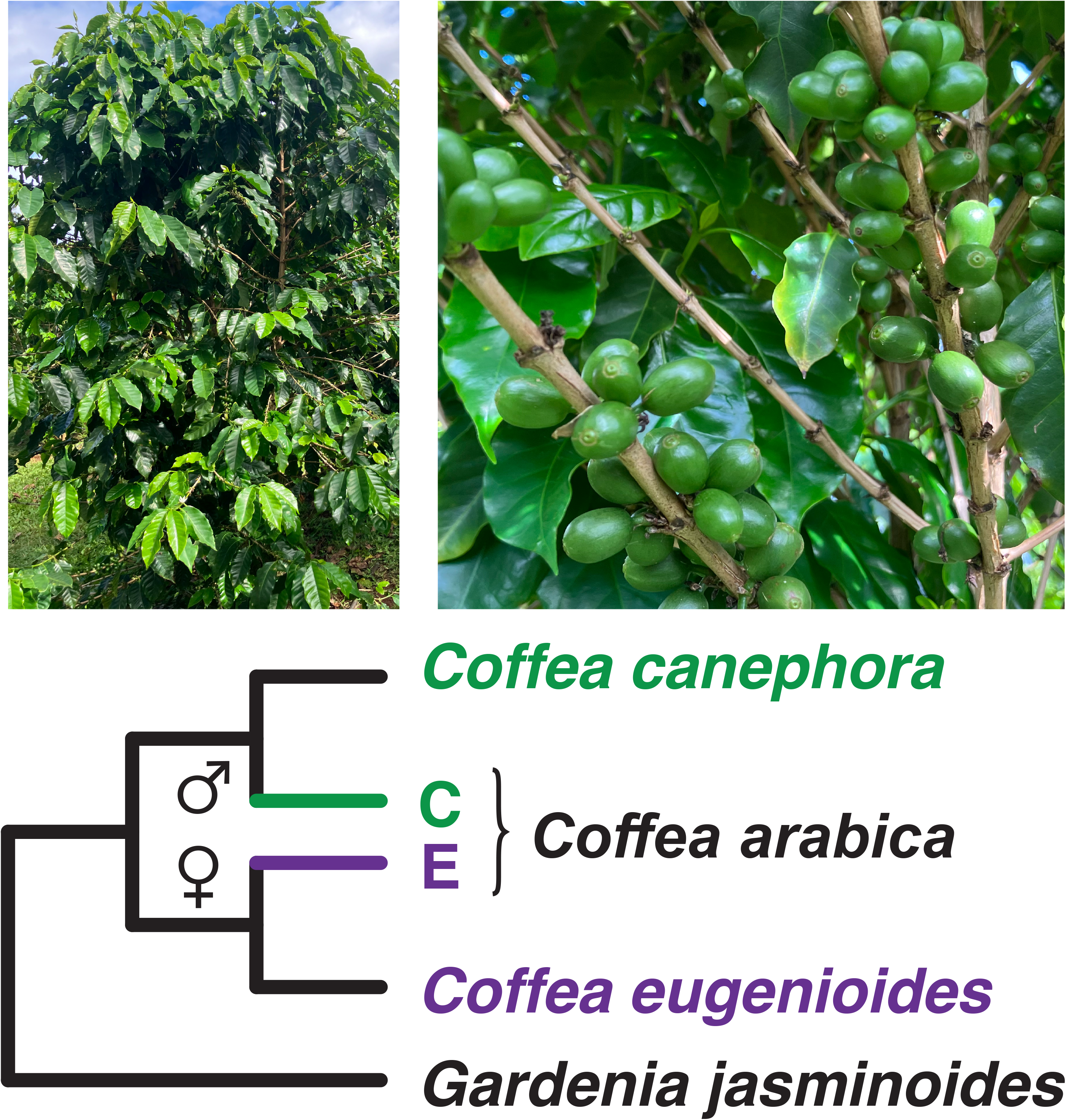
*Coffea arabica* is an allotetraploid formed via hybridization between *C. eugenioides* and *C. canephora*. Top: Images of *C. arabica* in a field in Oahu, HI. Image by A. Ortiz. Bottom: Cladogram depicting the relationships between *C. eugenioides*, *C. canephora*, hybrid tetraploid *C. arabica*, and outgroup *Gardenia jasminoides*. *Coffea eugenioides* served as the maternal (cytoplasmic) donor in the hybridization event, providing half of the nuclear genome and both organellar genomes (purple line), while *C. canephora* provided the paternal half of the nuclear genome (green line).

Here, we developed a new implementation of the classic ABBA-BABA test for inter-species introgression (Durand et al., 2011) to evaluate the direction and extent of HGF in allopolyploid genomes, and we employed it to characterize genome-wide patterns of HGF in *C. arabica*. To test whether HGF can contribute to the amelioration of epistatic incompatibilities in allopolyploids, we also evaluated whether genes whose products are targeted to the mitochondria and to the chloroplasts are especially likely to experience HGF from the maternally derived subgenome into the paternally derived subgenome, compared to genes whose products are targeted elsewhere in the cell. Overall, we found clear evidence of bi-directional HGF in the *C. arabica* genome, and evidence of maternally biased HGF in nuclear-encoded genes that function in plastid-nuclear enzyme complexes, but a dearth of HGF (in either direction) in nuclear-encoded genes whose products are subunits of mito-nuclear enzyme complexes. We also compared our method to a tree-based approach and found very similar patterns across both methods, indicating that the easier-to-implement *D*-statistic approach can be powerfully leveraged for detecting HGF in allopolyploids.

## METHODS

### Developing an ABBA-BABA test for homoeologous gene flow

A rooted, four-taxon bifurcating tree can have three possible topologies: 1) (((Sp1, Sp2), Sp3), O); 2) (((Sp1, Sp3), Sp2), O); and 3) (((Sp2, Sp3), Sp1), O). All three gene tree topologies will be observed if enough genes are sampled from the genome. In the absence of hybridization, the most common gene tree topology among these three possibilities will be identical to the species branching order, while the two rarer gene tree topologies are expected to be approximately equal in abundance, owing to the random fixation of polymorphic alleles across the two sequential splits (*i.e*., incomplete lineage sorting, ILS). Overabundance of one ILS tree compared to the other is a robust indicator of hybridization (Forsythe et al., 2020). The traditional ABBA-BABA test (Durand et al., 2011) takes advantage of this expectation to test whether nucleotide site patterns exhibit asymmetrical abundance across the genome (Figure S1). Assuming the true species tree of four species can be denoted as (((Sp1, Sp2), Sp3), O), the nucleotide site pattern in which the outgroup and Sp3 share an *ancestral* allele (A), whereas Sp1and Sp2 share a *derived* allele (B) is consistent with the species branching order. By contrast, a site pattern in which Sp1 and the outgroup share the ancestral allele (A) and Sp2 and Sp3 share the derived allele (B) represents one ILS tree topology (denoted an ABBA site pattern), and the opposite site pattern in which the ancestral allele is shared by Sp2 and outgroup and the derived allele is shared by Sp1 and Sp3 represents the other ILS tree topology (BABA site pattern). A relatively equal number of discordant ABBA and BABA site patterns is expected under ILS; however, an excess of either the ABBA or the BABA site pattern cannot be explained by ILS alone and requires some form of introgression between Sp2–Sp3 (overabundance of ABBA) or between Sp1–Sp3 (overabundance of BABA).

We extended this same logic to test whether HGF has impacted allopolyploid genomes. Specifically, we performed the ABBA-BABA test in two separate comparisons (Figure 2): 1) testing for HGF from the E genome into the C genome (*D_MAT_*), and 2) testing for HGF from the C genome into the E genome (*D_PAT_*). For each test, we quantified the number of sites for which derived alleles where shared by one subgenome and the opposing diploid genome (ABBA sites) compared to the number of sites for which derived alleles were shared by the two diploid genomes (BABA sites), using *Gardenia jasminoides* as an outgroup. If the number of ABBA sites was greater than the number of BABA sites (*i.e*., *D_MAT_* > 0 or *D_PAT_* > 0), we could infer HGF in that particular direction. In essence, this method tests whether the paternally derived subgenome has become more ‘maternal-like’, and whether the maternally derived subgenome has become more ‘paternal-like’, since the allopolyploidization event. We quantified *D* statistics for concatenated alignments of 6,672 genes, using 10,000 gene-level bootstrap replicates to assess statistical significance (Python scripts are available at https://github.com/albuquerque-turkey/Coffea_HGF). We calculated 95% CIs from the gene-level bootstrap replicates and performed a Z-test to determine whether *D* statistics departed significantly from 0. *D*-statistic point estimates, bootstrap distributions, and 95% CIs were plotted in R v4.1.2 (Team and Core Team, n.d.) using *ggplot2* (Wickham, 2011).

**Figure 2.**
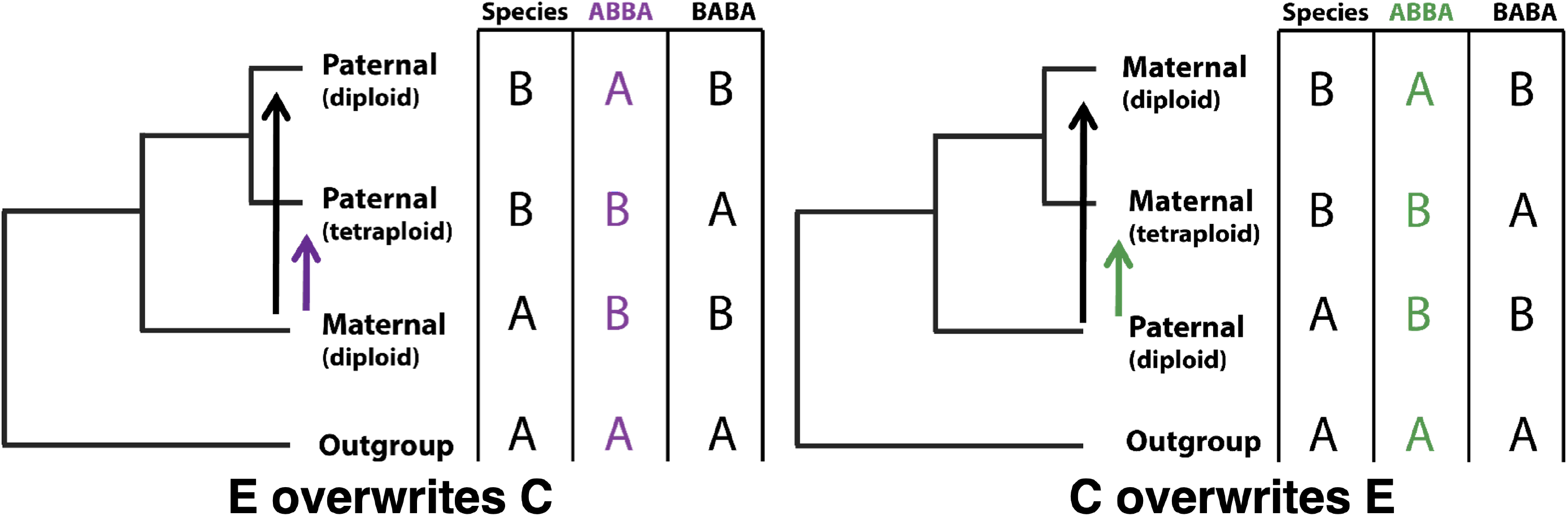
Two-way, reciprocal ABBA-BABA test on *C. arabica* subgenomes. The two-way reciprocal ABBA-BABA test leverages the sister relationship between each subgenome and its corresponding diploid to estimate the rate of HGF from the other subgenome (left – maternal overwrites paternal, right – paternal overwrites maternal). The “donor” subgenome is excluded from the test, such that HGF between subgenomes is expected to produce an “ABBA” site pattern, with subgenomes appearing to be more closely related to the opposite diploid than to the diploid from which they were originally derived. The opposite “BABA” site pattern is therefore used as an estimate of the expected frequency of this pattern evolving by random chance (*i.e*., ILS).

### Orthologous gene alignments and CyMIRA gene classification

Our full HGF analysis was run on 6,672 orthologous single-copy genes (Table S1), originally identified in (Sharbrough et al., 2022), with alignments available at doi.org/10.6084/m9.figshare.24085830. Briefly, we obtained CDS sequences from publicly available assemblies for *C. arabica* (GCF_003713225.1)*, C. eugenioides* (GCF_003713205.1)*, C. canephora* (GCA_900059795.1), and *G. jasminoides* (GCA_013103745.1), aligned all 6,672 orthologous gene groups with MAFFT v7.480 (Katoh and Standley, 2013), using a perl wrapper (https://github.com/dbsloan/perl_modules/sloan.pm) to convert the CDS sequences to amino acid sequences, align with MAFFT, and then convert the sequences back to nucleotides, as in (Sharbrough et al., 2022). We used two distinct alignment trimming strategies: Gblocks v0.91b with the ‘-n’ parameter set (Castresana, 2000), and ClipKIT v1.2.0 (Steenwyk et al., 2020) with the −l parameter set and using a custom Python wrapper to convert ClipKIT-trimmed amino acid alignments back to CDS alignments (https://github.com/jsharbrough/protTrim2CDS). Gblocks-trimmed and ClipKIT-trimmed alignments that were ≥ 99bp in length were fed into ABBA-BABA tests to ensure that trimming method did affect our inferences of HGF.

To evaluate whether HGF was especially maternally biased in genes targeted to, and interacting with, mitochondrial and chloroplast genes and gene products, we used the classifications from Sharbrough et al., (2022), available at https://github.com/jsharbrough/CyMIRA_gene_classification/tree/master/Species_CyMIRA to classify genes into six categories: non-organelle-targeted (NOT), mitochondrial-targeted, non-interacting (MTNI), mitochondrial-targeted, interacting (MTI), mitochondrial enzyme complexes (MTEC), plastid-targeted, non-interacting (PTNI), plastid-targeted, interacting (PTI), and plastid enzyme complexes (PTEC) (Table 1). Of those, the non-interacting and interacting categories were mutually exclusive for each organelle, while genes involved in enzyme complexes were a subset of interacting genes that are involved in cytonuclear enzyme complexes, as defined by CyMIRA v1.0 (Forsythe et al., 2019). Mitochondrial enzyme complexes included the mitochondrial TAT complex, the mitochondrial ribosome, NADH dehydrogenase (Complex I of the oxidative phosphorylation cascade – OXPHOS), ubiquinol-cytochrome c reductase (Complex III of OXPHOS), cytochrome c oxidase (Complex IV of OXPHOS), and ATP synthase (Complex V of OXPHOS). Plastid enzyme complexes included the heteromeric ACCase, the chloroplast ribosome, the CLP protease, both photosystems I and II, Rubisco, the chloroplast NADH dehydrogenase-like complex, cytochrome b6f, and the chloroplast ATP synthase. Gene names for each of these complexes from all five species are available in Table S2.

**Table 1.**
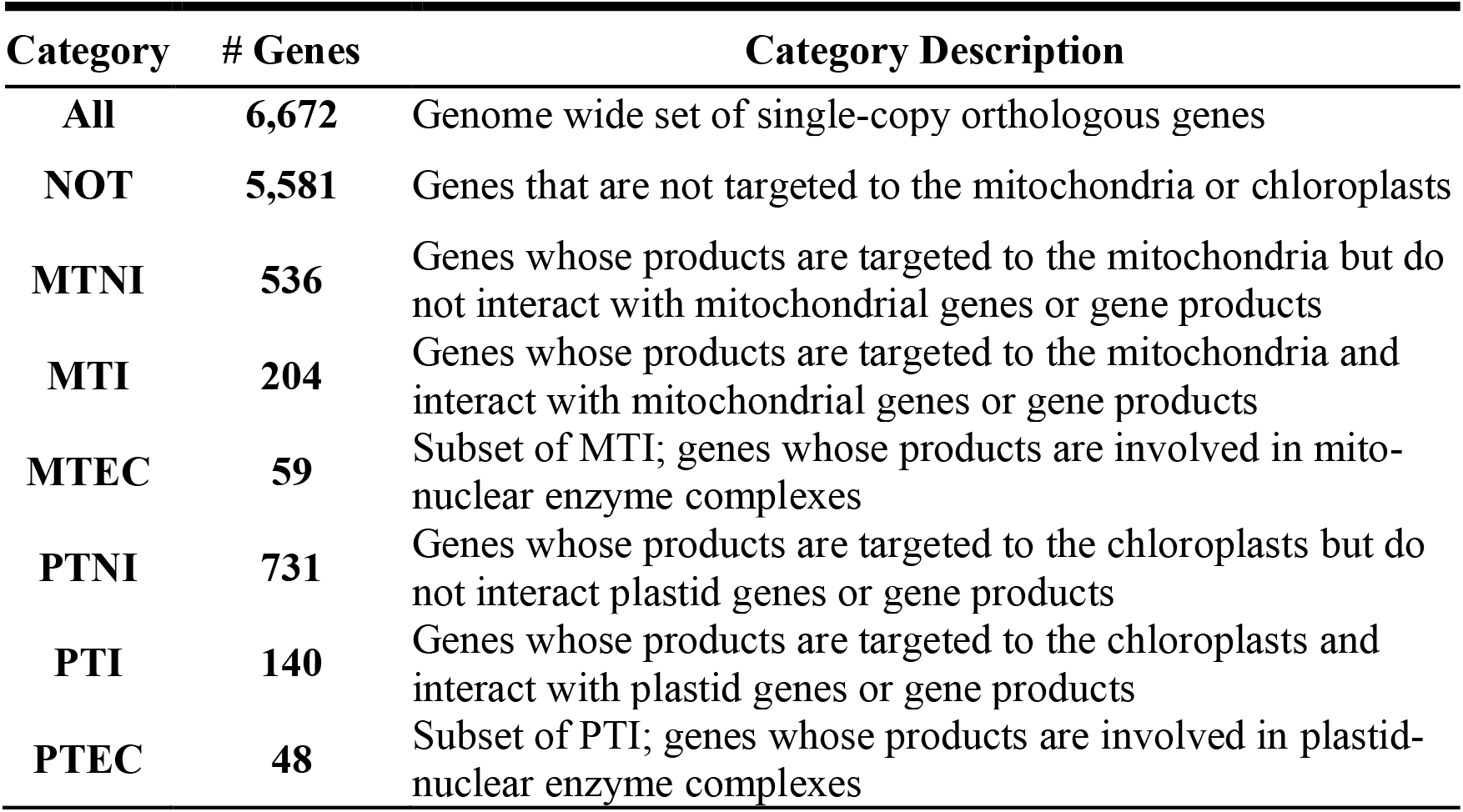
Functional classification of single-copy orthologous genes in *Coffea*.

### Phylogenetic analyses

To validate the inferences made by our newly developed method, we inferred phylogenetic trees by Maximum Likelihood using RAxML v8.2.12 (Stamatakis, 2014). For each gene tree, we used the raxmlHPC-PTHREADS function, employing the rapid bootstrap analysis and search for best-scoring ML tree in one program run (‘-f a’ parameter), with 100 bootstrap replicates, and assuming the GTRGAMMAIX model of molecular evolution. We rooted each of the resulting gene trees with the *G. jasminoides* sequence and determined which of the 15 possible gene tree topologies (Figure S2) each gene tree fit. We reasoned that if HGF was unidirectional in a given gene, we expected tetraploids to be sister to each other, with the direction of HGF matching the most closely related diploid (tree topology L or M – Figure S3). Of course, these topological patterns can also be produced by ILS, recurrent mutations, and autapomorphies; however, in the absence of HGF, the probability of the tetraploids to be sister to each other should be equal to the probability of a tetraploid being sister to the opposing diploid (tree topology N or H) and to that of the two diploids being sister to on another (tree topology I or O). Similarly, if HGF resulted in a reciprocal exchange of DNA, we would observe a tree in which tetraploid sequences would group with the opposing subgenome (tree topology F), while trees derived from random processes would find the diploids as sister to one another and the tetraploids as sister to one another (tree topology K). The relative abundance of the HGF trees compared to their ILS-derived alternatives provides a similar comparison to the *D*-statistic, but based on whole genes and an explicit model of molecular evolution. We compared HGF tree topology abundance using a series of binomial, χ^2^, and Fisher’s Exact tests, correcting for multiple comparisons using the Holm procedure for the Bonferroni correction (Holm, 1979).

We also tested whether putative HGF trees were especially abundant among mitochondrial and plastid enzyme complex genes, using a Fisher’s Exact test to determine whether cytonuclear enzyme complex genes exhibited an increase in the ratio of maternally biased HGF trees compared to non-organelle-targeted genes.

## RESULTS

*Testing for HGF in allotetraploid* C. arabica

We used a stepwise, reciprocal ABBA-BABA framework to evaluate whether homoeologous exchange and homoeologous gene conversion contribute to gene flow between subgenomes of allotetraploid *C. arabica*. These analyses of the ClipKIT-trimmed alignments of 6,672 orthologous genes revealed strong evidence of bidirectional HGF (*i.e*., *D_MAT_* and *D_PAT_* >> 0; Table 2), as we saw extensive overwriting of the C subgenome by the E subgenome (maternally derived HGF) and overwriting of the E subgenome by the C subgenome (paternally derived HGF) (Figure 3). Similar results were obtained from Gblocks-trimmed alignments, which are provided in the Supplementary material (Figure S4, Table S3). Maternally derived HGF (*D_MAT_* (CI_95_) = 0.538 (0.448 – 0.610)) was slightly, but not significantly greater than paternally derived HGF after correcting for multiple comparisons (*D_PAT_* (CI_95_) = 0.416 (0.357-0.475); Z = 2.785; *p* = 0.0026). Overall, our new method indicated the presence of substantial and bidirectional HGF in *C. arabica* genomes.

**Figure 3.**
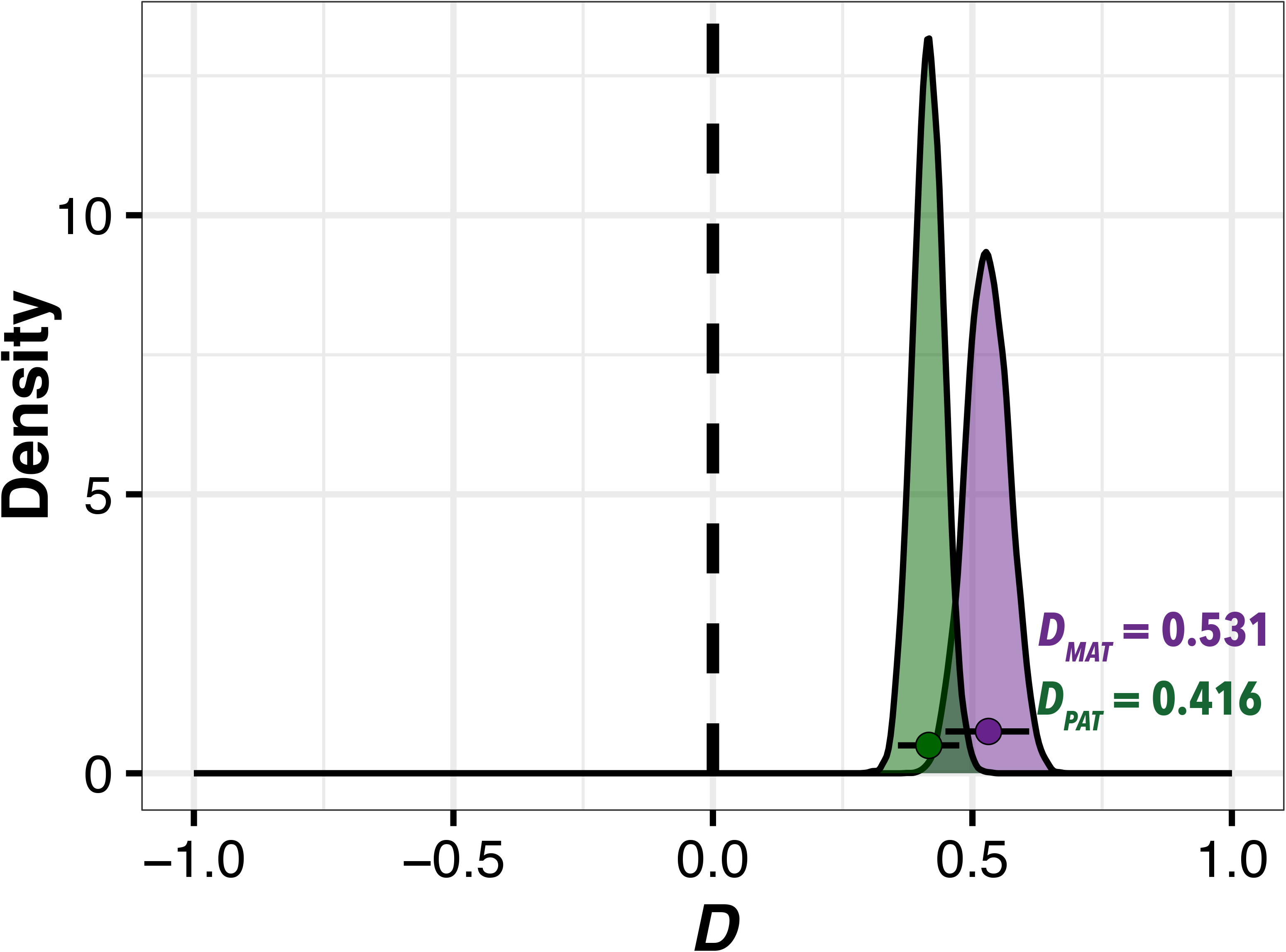
Genome-wide patterns of bidirectional HGF in allotetraploid *C. arabica*. Reciprocal *D*-statistic estimates of the E subgenome overwriting the C subgenome (purple, *Dmat*) and of the C subgenome overwriting the E subgenome (green, *Dpat*). Points represent overall *D*-statistics, density plots depict distributions from 10,000 gene-level bootstrap replicates, and error bars represent 95% CIs. Distributions that are significantly greater than 0 are indicative of HGF in that direction.

**Table 2.**
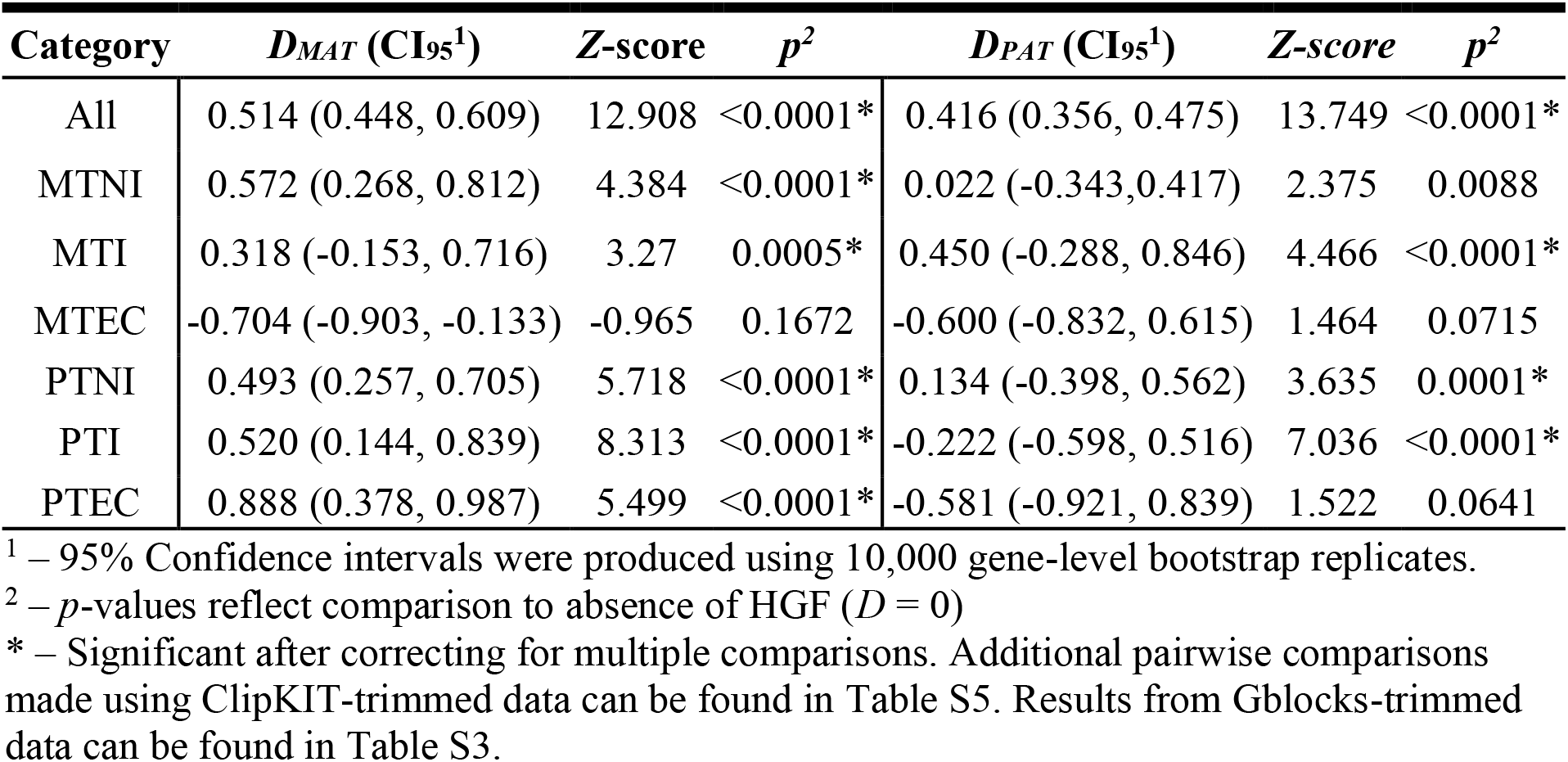
Reciprocal ABBA-BABA test statistics across gene functional categories.

To evaluate the robustness of these results, we also inferred phylogenetic trees for each of these 6,672 genes. Overall, the majority of genes exhibited the same topology as the species tree (3,671/6,672, 54.7%); however, a sizeable fraction of genes exhibited alternative topologies, and all 15 possible tree topologies were observed in these genes (Table 3, Figure 4)). Among these, there were 1,115 genes (16.7%) in which the E subgenome, the C subgenome, and *C. eugenioides* formed a clade to the exclusion of *C. canephora.* Of these, 316 genes (4.7%) exhibited sister relationships between the E and C subgenomes. We denote these 316 trees as putative E overwriting C trees. Similarly, there were 1,221 genes (18.3%) in which the C subgenome, the E subgenome, and *C. canephora* all formed a clade to the exclusion of *C. eugenioides*, with 289 genes (4.3%) in which the subgenomes were sister to one another (putative C overwriting E trees). There were also 198 genes (3.0%) in which the C subgenome grouped with *C. eugenioides* and the E subgenome grouped with *C. canephora* (putative reciprocal exchange trees). Together, these phylogenetic results provide a very similar picture as the reciprocal ABBA-BABA test, with a substantial fraction of the *C. arabica* exhibiting patterns consistent with HGF.

**Figure 4.**
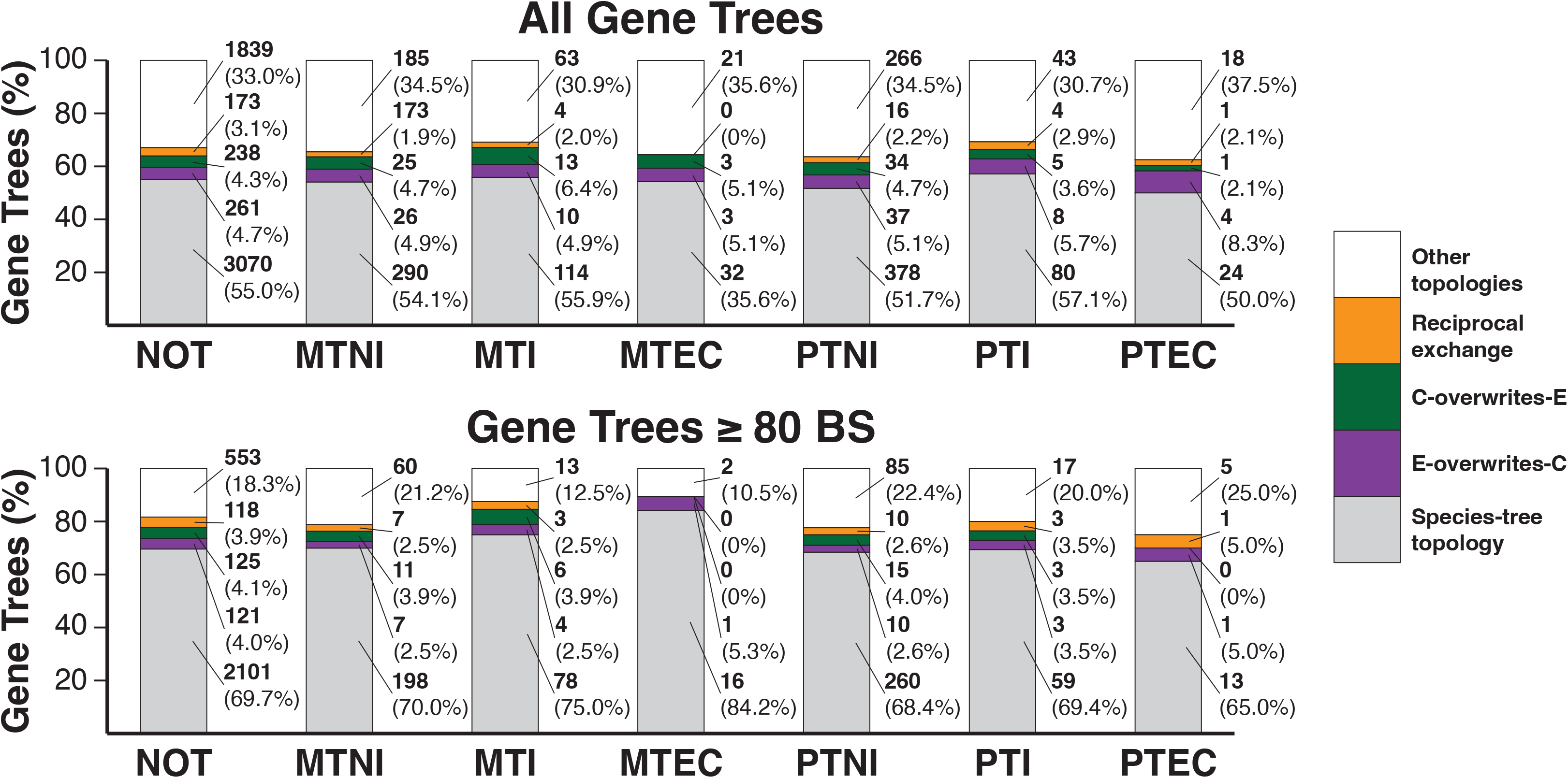
Single-copy orthologous gene tree topologies in *Coffea*. Percentage of gene trees (top: all trees, bottom: only trees with >80 bootstrap support) of various tree topologies (species topology – gray; E-overwriting-C – purple; C-overwriting-E – green; reciprocal exchange – orange; other – white) in single-copy orthologous gene groups.

**Table 3.** Distribution of gene tree topologies across functional gene categories.

| **Topology<sup>1</sup>** | **ALL** | **NOT** | **MTNI** | **MTI** | **MTEC** | **PTNI** | **PTI** | **PTEC** |
| --- | --- | --- | --- | --- | --- | --- | --- | --- |
| A<sup>2</sup> | 3651 | 3070 | 290 | 114 | 32 | 378 | 80 | 24 |
| B | 478 | 410 | 30 | 10 | 1 | 49 | 8 | 1 |
| C | 278 | 230 | 24 | 8 | 7 | 36 | 4 | 2 |
| D | 474 | 396 | 39 | 12 | 2 | 55 | 9 | 5 |
| E | 629 | 507 | 67 | 18 | 4 | 88 | 11 | 3 |
| F<sup>3</sup> | 198 | 173 | 10 | 4 | 0 | 16 | 4 | 1 |
| G | 89 | 75 | 6 | 2 | 2 | 8 | 4 | 3 |
| H | 34 | 31 | 2 | 0 | 0 | 3 | 0 | 0 |
| I | 19 | 13 | 2 | 2 | 0 | 4 | 0 | 0 |
| J | 68 | 57 | 6 | 2 | 0 | 7 | 3 | 1 |
| K | 81 | 63 | 8 | 5 | 1 | 11 | 2 | 1 |
| L<sup>4</sup> | 316 | 261 | 26 | 10 | 3 | 37 | 8 | 4 |
| M<sup>5</sup> | 289 | 238 | 25 | 13 | 3 | 34 | 5 | 1 |
| N | 37 | 31 | 0 | 2 | 2 | 2 | 2 | 2 |
| O | 31 | 26 | 1 | 2 | 2 | 3 | 0 | 0 |
| **Total** | **6672** | **5581** | **536** | **204** | **59** | **731** | **140** | **48** |

<sup>1</sup> – See Figure S2 for all gene tree topologies<sup>2</sup> – Species tree<sup>3</sup> – Tree consistent with reciprocal exchange across subgenomes<sup>4</sup> – Tree consistent with C subgenome overwriting E subgenome<sup>5</sup> – Tree consistent with E subgenome overwriting C subgenome

To evaluate the likelihood that these trees arose through ILS, we compared the number of putative E overwriting C trees, the number of putative C overwriting E trees, and the number of putative reciprocal exchange trees to the number of other trees that would be expected by ILS (Figure S3). We found significantly higher numbers in all three sets of putative HGF gene trees compared to their ILS counterparts (Table S4), a pattern that was even true when restricting the analysis to those trees with ≥80 bootstrap support. Together, the prevalence of putative HGF trees is unlikely to be explained entirely by ILS, indicating that HGF has played a prominent role in the genealogical history of the *C. arabica* genome.

*Maternally biased HGF in plastid-targeted, but not mitochondrial-targeted nuclear genes Coffea arabica* received both its mitochondrial and plastid genomes from *C. eugenioides*, but nuclear DNA from both *C. eugenioides* and *C. canephora* (Cros et al., 1998). Because interactions between nuclear-encoded genes and mitochondrial- and plastid-encoded genes are critical for plant function and fitness (Kremnev and Strand, 2014; Kühn et al., 2015), mutational changes in one genome or the other are expected to produce intense selection for compensatory changes to maintain respiratory and photosynthetic function (Rand et al., 2004). It therefore stands to reason that co-adapted mito-nuclear and plastid-nuclear epistatic interactions might be disrupted upon genome merger (Sharbrough et al., 2017). If conflict between the paternally derived nuclear subgenome and the maternally derived cytoplasmic genomes exists, HGF provides a rapid evolutionary mechanism to ameliorate the cytonuclear mismatch by replacing paternally derived genes targeted to the energy-producing organelles with maternally derived genes. We tested this hypothesis using our reciprocal ABBA-BABA method in sets of organelle-targeted non-interacting genes, interacting genes, and genes involved in cytonuclear enzyme complexes. There was no apparent difference between genome-wide patterns of HGF and organelle-targeted genes that do not interact with organelle genes or gene products (n_MTNI_ = 536; n_PTNI_ = 731; Figure 5a and 5b). Interacting genes (n_MTI_ = 204; n_PTI_ = 140) also showed a similar pattern (Figure 5c and 5d), but we found substantially different patterns of HGF compared to non-organelle-targeted genes in the subset of interacting genes that are involved in cytonuclear enzyme complexes (n_MTEC_ = 59; n_PTEC_ = 48). Mitochondrial enzyme complex genes exhibited no evidence of HGF in either direction (*D_MAT_* (CI_95_) = −0.255 (−0.692, 0.345); Z = −0.965; *p* = 0.167; *D_PAT_* (CI_95_) = 0.333 (−0.200, 0.684); Z = 1.464; *p* = 0.0715) (Figure 5e), indicating that HGF is not acting to ameliorate mitonuclear conflicts in *C. arabica*. By contrast, plastid enzyme complex genes exhibited substantial amounts of maternally biased HGF (*D_MAT_* (CI_95_) = 0.750 (0.415, 0.944); Z = 5.499; *p <* 0.0001), but no evidence of paternally biased HGF (*D_PAT_* (CI_95_) = 0.500 (−0.333, 0.916); Z = 1.522; *p* = 0.0641) (Figure 5f, Table S5). This pattern of the E subgenome overwriting the C subgenome in this set of genes is consistent with a scenario in which cytonuclear mismatches caused by the paternally derived nuclear subgenome are ameliorated by maternally biased HGF.

**Figure 5.**
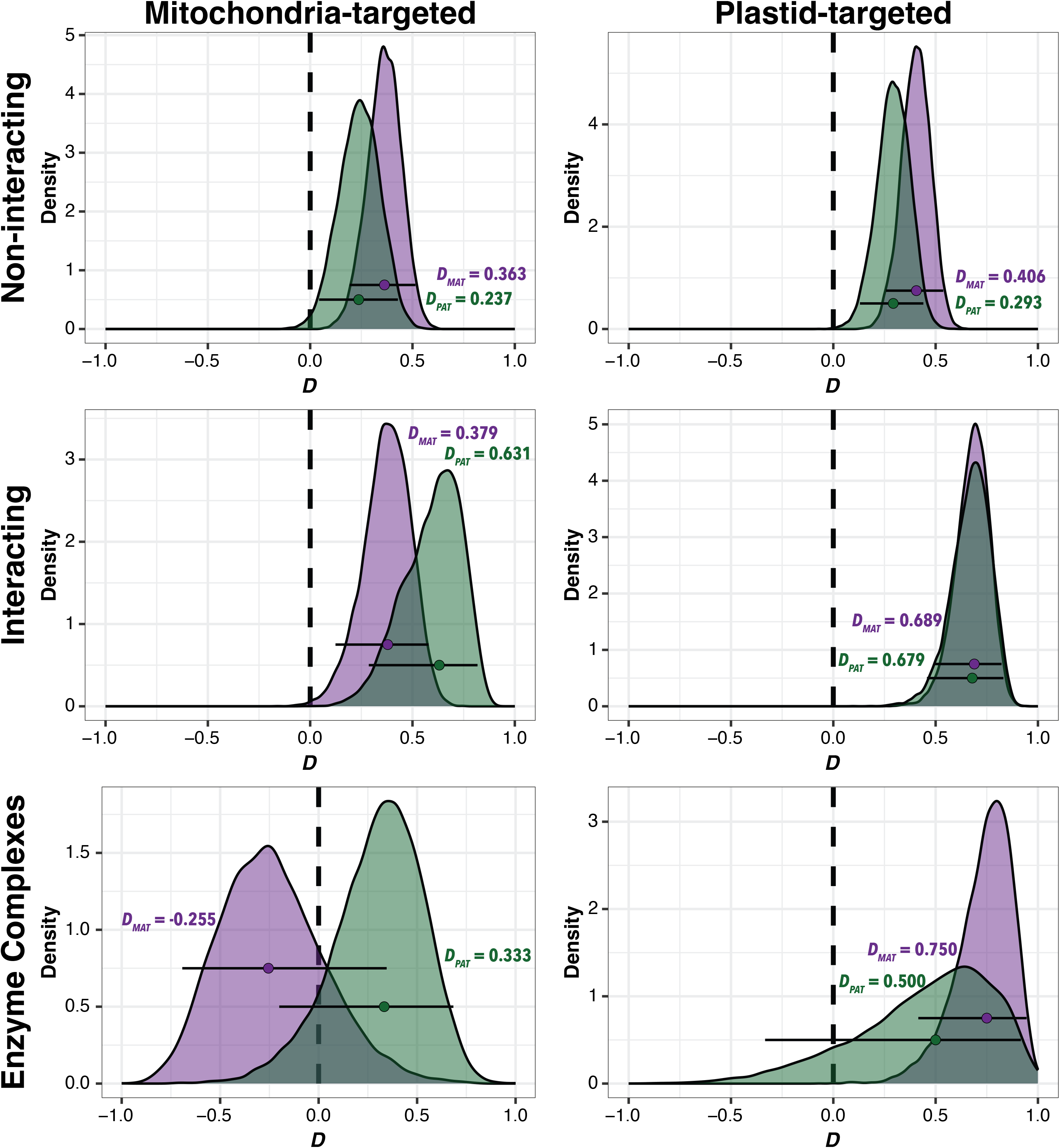
Patterns of HGF in mitochondria- and plastid-targeted genes in allotetraploid *C. arabica*. Reciprocal *D*-statistic estimates of the E subgenome overwriting the C subgenome (purple, *Dmat*) and of the C subgenome overwriting the E subgenome (green, *Dpat*) in genes targeted to the mitochondria (left) and to the plastid (right). Genes are grouped according to the intimacy of interaction: non-interacting – top panels, interacting – middle panels, enzyme complexes – bottom panels. Points represent overall *D*-statistics, density plots depict distributions from 10,000 gene-level bootstrap replicates, and error bars represent 95% CIs. Distributions that are significantly greater than 0 are indicative of HGF in that direction.

We also tested whether cytonuclear enzyme complex genes exhibited evidence of maternally biased HGF based on gene trees. We observed a higher proportion of plastid enzyme complex genes exhibited the E-overwriting-C topology (n = 4, 8.3%) and a lower proportion of plastid enzyme complex genes exhibited the C-overwriting-E topology (n = 1, 2.1%) compared to the non-organelle-targeted genes (n_E-over-C_ = 261, 4.7%; n_C-over-E_ = 238, 4.3%), but neither difference was significant (Fisher’s Exact Test, *p*_E-over-C_ = 0.274; *p*_C-over-E_ = 1.0). Notably, the genes that did exhibit maternally biased HGF topologies are all central members of four distinct and essential plastid enzyme complexes: Rubisco, NDH, PSI, and CLP (Figure 6). An equal number of mitochondrial enzyme complex genes exhibited the E-overwriting-C topology as the C-overwriting-E topology (n = 3, 5.1% for each), which were not different from the pattern in non-organelle-targeted genes (*p*_E-over-C_ = 0.752; *p*_C-over-E_ = 0.737). Although the patterns of HGF in plastid-targeted enzyme complex genes were not significantly different from the genome-wide pattern, this change in proportion is in the direction we predicted (and is consistent with our reciprocal ABBA-BABA results), potentially indicating that the phylogenetic approach lacks the power to reveal biased HGF as effectively as our newly implemented method.

**Figure 6.**
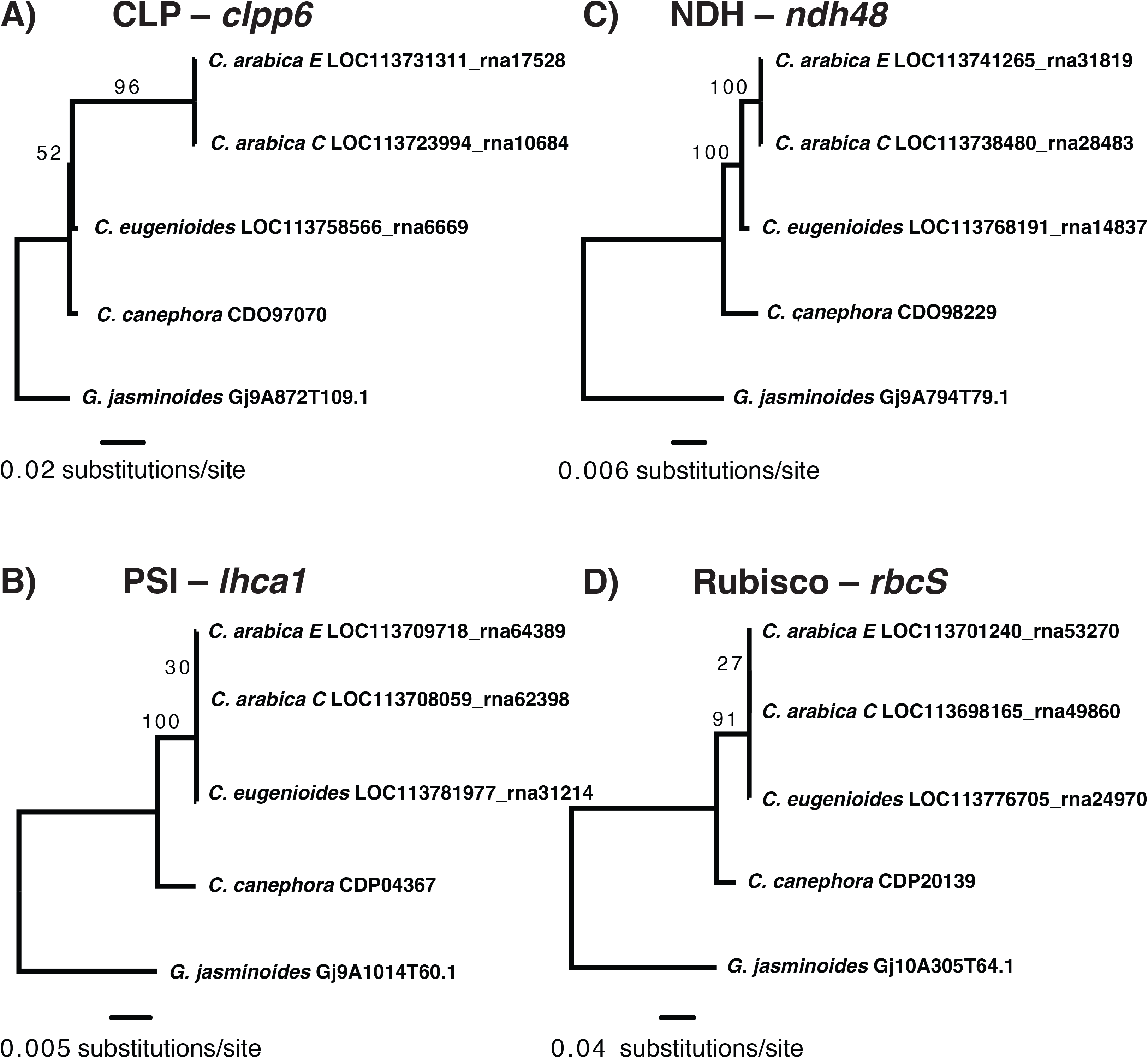
Gene trees of four plastid enzyme complex genes that appear to exhibit maternally biased HGF. A) Gene tree for *clpp6*, a nuclear-encoded gene whose protein product participates in the heteromeric ring of the CLP protease. B) Gene tree for *ndh48*, a nuclear-encoded gene whose protein product is involved in the plastid NDH. C) Gene tree for *lcha1*, a nuclear-encoded gene whose protein product is involved in photosystem I. D) Gene tree for *rbcS*, a nuclear-encoded gene whose protein product represents the small subunit of the Rubisco protein complex.

## DISCUSSION

### A stepwise, reciprocal ABBA-BABA test to detect HGF in allopolyploid genomes

Recombination between homoeologs provides an important source of novel variation for allopolyploids (Mason and Wendel, 2020), and characterizing patterns and consequences of HGF represents an important endeavor for understanding evolutionary dynamics after WGDs (Deb et al., 2023). This can be done in a number of ways (Schiessl et al., 2019) including comparative read mapping with long reads (*e.g*., (Jarvis et al., 2017; Chu et al., 2021; Oruganti et al., 2023)), single-nucleotide variant comparisons (*e.g.*, (Salmon et al., 2010; Lashermes et al., 2016)), structural variant characterization (*e.g*., (Orantes-Bonilla et al., 2022)), and even cytogenetic tools like fluorescent *in situ* hybridization (*e.g.*, (Stein et al., 2017)) or optical mapping (*e.g.,* (Yuan et al., 2018)). The method we implemented here incorporates the logic developed by the introgression literature (recently reviewed by (Hibbins and Hahn, 2022)), to evaluate HGF against an ILS framework. The ABBA-BABA statistic has been used to test for the presence of interspecies introgression in polyploid genomes before (*e.g*., (Pont et al., 2019)), but here we repurposed it to characterize patterns of introgression *within the same cell*.

Our method can also look for patterns of HGF in functionally related genes (*e.g*., genes whose products are targeted to the mitochondria and plastids). Because these genes are dispersed across the genome and not closely linked, it is possible to infer the activity of natural selection acting to fix HGF in a particular direction. Our current implementation is powerful enough to detect (or not detect) HGF with as few as ∼50 genes, but smaller gene sets could likely be tested by bootstrapping at the site level, rather than at the gene level.

Importantly, this reciprocal ABBA-BABA approach requires that introgression between diploid relatives not be extensive, as that would increase the abundance of BABA sites in our test and obscure patterns of HGF. Therefore, we recommend testing for diploid introgression prior to testing for HGF (we found no evidence of introgression between *C. eugenioides* and *C. canephora* here). This approach also relies upon high quality genome assemblies of the polyploid subgenomes, the two diploid relatives, and of an outgroup. Due to the availability of Pacbio HiFi reads, Hi-C (or similar) sequencing, and ultra-long Oxford Nanopore reads, among other technologies, such genomic resources are becoming increasingly available. The coffee genomes used in the present study may represent an important source of error, as they are extremely gappy (*C. arabica* assembly GCF_003713225.1 is comprised of 3,522 separate contigs), such that misassemblies could contribute to overestimating the rate of HGF. Continuing to improve genome assemblies (and producing pangenomes) will greatly facilitate our ability to evaluate the patterns and consequences of HGF in allopolyploids.

### Patterns of HGF in organelle-targeted genes

Hybrid incompatibilities between nuclear and cytoplasmic genomes can be produced via cytonuclear co-evolution, and this special class of BDMIs can play a prominent role in reinforcing species boundaries (Burton and Barreto, 2012; Sloan et al., 2017). Polyploidy may offer a general solution to cytonuclear incompatibilities, as the maternally derived nuclear genes are retained following the genome merger event. As a result, those genes can act as sources of epistatic variation co-adapted with the cytoplasmic genomes. This is expected to result in either the loss of paternally derived genes encoding products targeted to the mitochondria or chloroplasts, HGF that overwrites paternally derived homoeologs with maternally derived copies, or rapid evolution in paternally derived copies under selection to more closely “match” cytoplasmic interacting partners (Sharbrough et al., 2017).

In a broad sampling of angiosperm allotetraploids, Sharbrough and colleagues (2022) demonstrated that paternally derived organelle-targeted genes do not experience global accelerations in rate, and are not lost at higher rates than maternally derived copies (Sharbrough et al., 2022). A number of allopolyploid angiosperms exhibit evidence of maternally biased homoeologous gene conversion of *rbcS*, the nuclear-encoded subunit of Rubisco (Gong et al., 2012, 2014; Li et al., 2020). Here, we found that paternally derived genes whose products are involved in jointly encoded plastid-nuclear enzyme complexes are preferentially overwritten by maternally derived genes, but no such pattern was observed in paternally derived genes involved in mitochondrial-nuclear enzyme complexes.

There are several potential explanations for the discrepancy across organellar compartments. First, plastid genomes typically evolve at much higher rates than mitochondrial genomes in angiosperms (Wolfe et al., 1987), and cytonuclear incompatibilities are unlikely to occur in the absence of accelerated rates of cytoplasmic genome evolution (Havird et al., 2015; Rockenbach et al., 2016; Williams et al., 2019). Indeed, the apparent lack of HGF in either direction in mitochondrial enzyme complex genes that we observed here may reflect an absence of mitonuclear incompatibilities in *C. arabica*, owing to a low rate of mitochondrial genome evolution and low sequence divergence between mitochondrial genomes of *C. eugenioides* and *C. canephora*. Second, the efficacy of selection acting on plastid-nuclear interactions may be greater than that acting on mitochondrial-nuclear interactions. Plastid enzyme complexes are expressed at much higher rates than mitochondrial enzyme complexes (Forsythe et al., 2022), to the extent that Rubisco is likely the most abundant enzyme on Earth (Ellis, 1979; Raven, 2013). Genes expressed at higher levels experience more effective selection than genes expressed at lower levels (Drummond et al., 2005; Yang et al., 2012). Additionally, plastid genomes are retained at copy numbers orders of magnitude higher than mitochondrial genomes (Fernandes Gyorfy et al., 2021), which is expected to produce a higher effective population size in plastid genomes than mitochondrial genomes.

### Summary & Conclusions

Here, we implemented a repurposing of an existing method for detecting introgression between species into a method for detecting gene flow within a cell and tested this method in the allotetraploid angiosperm *C. arabica*. This proof-of-principle for a simple method for detecting homoeologous gene flow in allopolyploid taxa provides an important tool for characterizing the impacts of homoeologous exchange and gene conversion in allopolyploids. We also document patterns of maternally biased HGF in plastid-targeted, but not mitochondria-targeted genes that are involved in plastid-nuclear enzyme complexes.

## ACKNOWLEDGEMENTS

This work was supported by grants from the National Science Foundation Plant Genomes Research Program (awards 1829176 and 2145811). Primary analyses were performed on the *Logrus* HPC operated by the National Center for Genome Resources, Santa Fe, New Mexico, and later analyses were performed on Alpine, which is a jointly funded HPC by the University of Colorado Boulder, the University of Colorado Anschutz, Colorado State University, and the National Science Foundation (award 2201538).

## AUTHOR CONTRIBUTIONS

AO wrote the code for the method, completed all analyses presented here, and wrote the first draft of the manuscript. JS conceived of the method, assisted with validating code, and edited the manuscript. Both authors contributed equally to figure and table development.

## DATA AVAILABILITY

All genomic data used in this data are already publicly available. Alignments and trees for orthologous gene groups are available at doi.org/10.6084/m9.figshare.24085830. All code developed for this project is available at https://github.com/albuquerque-turkey/Coffea_HGF.

---

**Figure S1.**
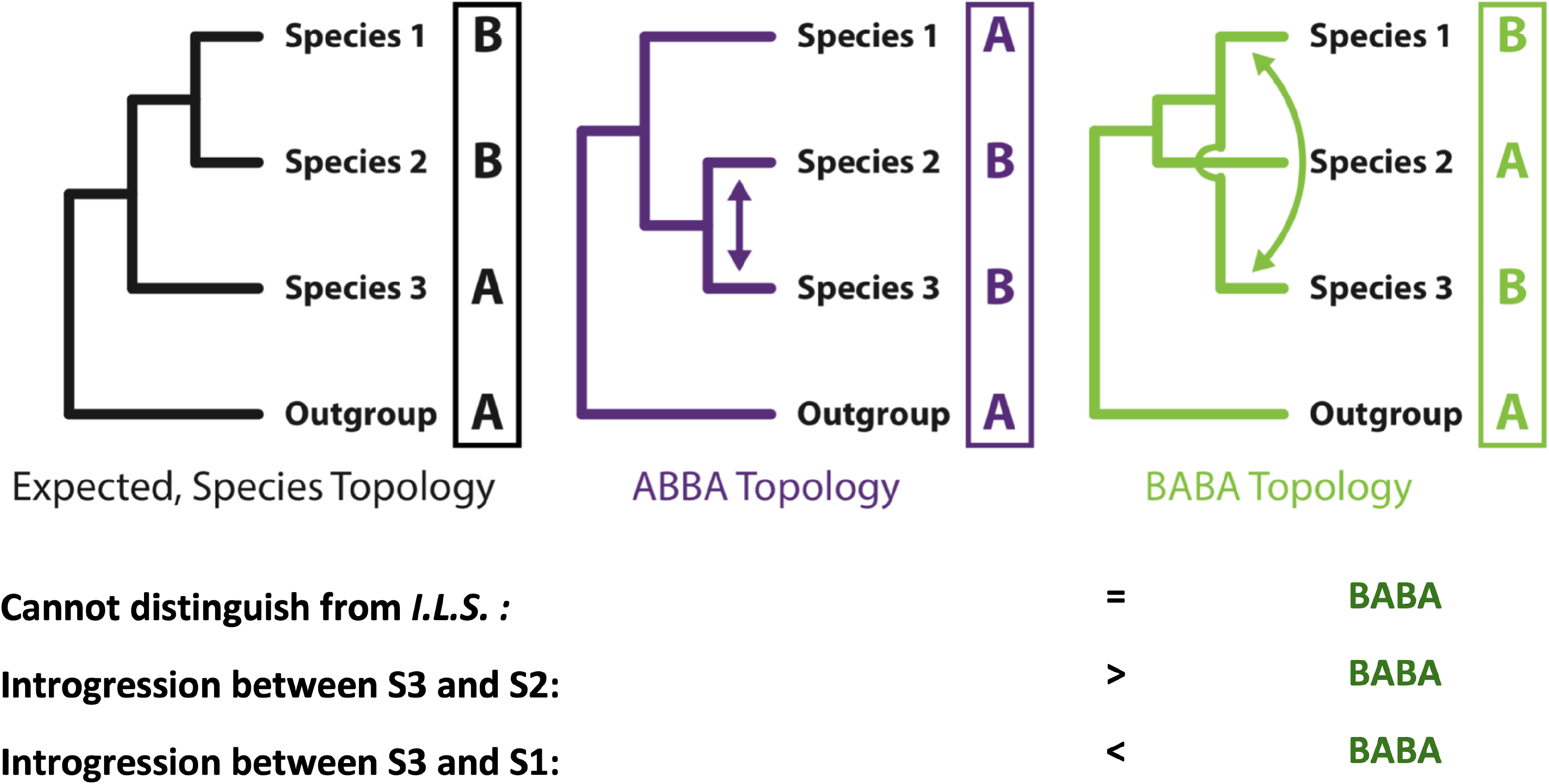
Classic implementation of the ABBA-BABA test. The relative abundance of ABBA (middle tree) vs. BABA (right tree) site patterns in genome-wide alignments can be used to infer introgression between species in a four-taxon arrangement.

**Figure S2.**
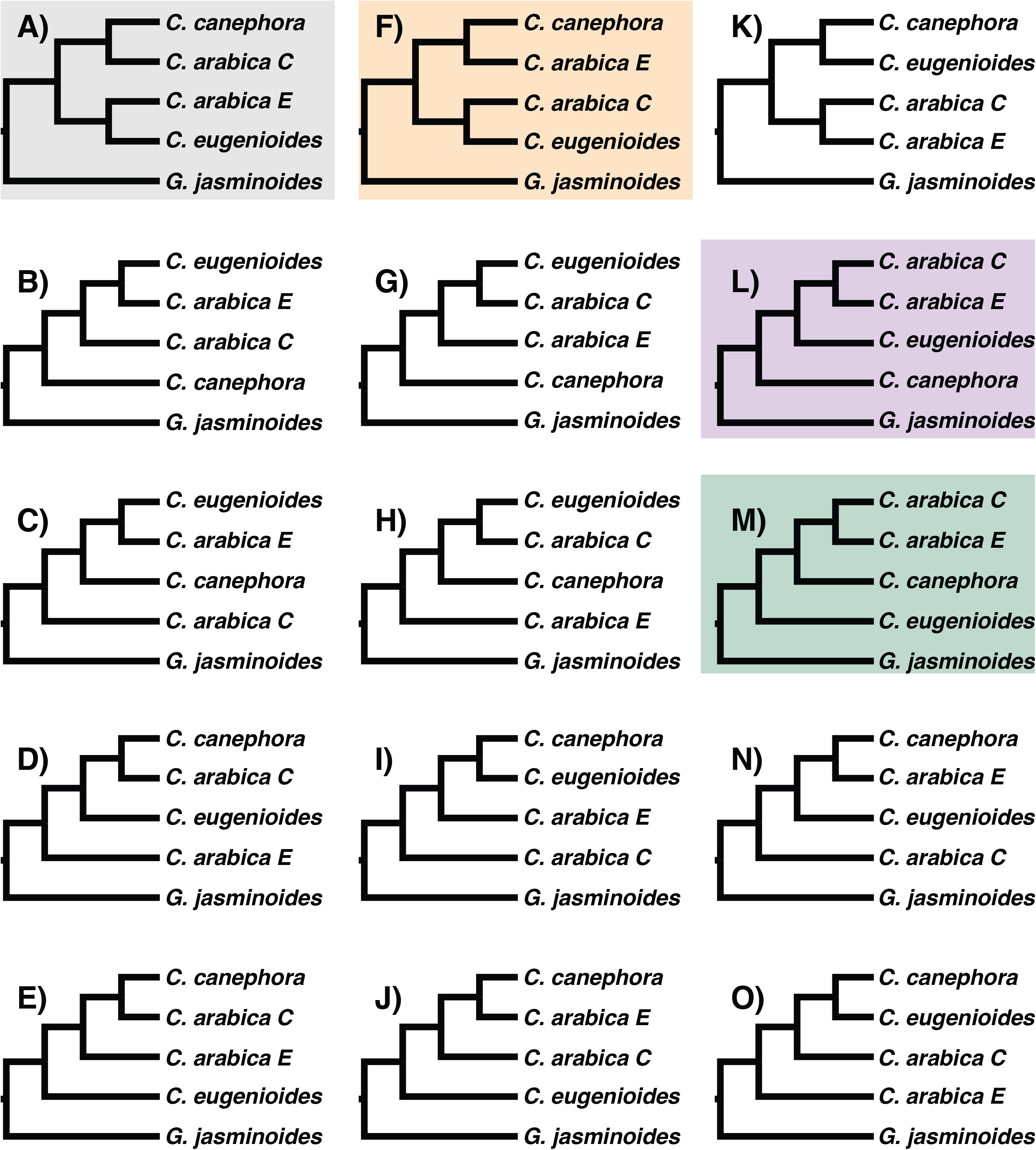
Gene tree topologies for rooted, five-taxon trees. There are 15 possible tree topologies for a rooted tree with five taxa. The *Coffea* species tree is depicted in A), and is highlighted in gray. The gene tree that would be expected if HGF were reciprocal across subgenomes is depicted in F) (highlighted in orange). The gene tree that would be expected if HGF were maternally biased (*i.e.*, E-overwriting-C) is depicted in L) (highlighted in purple). The gene tree that would be expected if HGF were paternally biased (*i.e.*, C-overwriting-E) is depicted in M) (highlighted in green).

**Figure S3.**
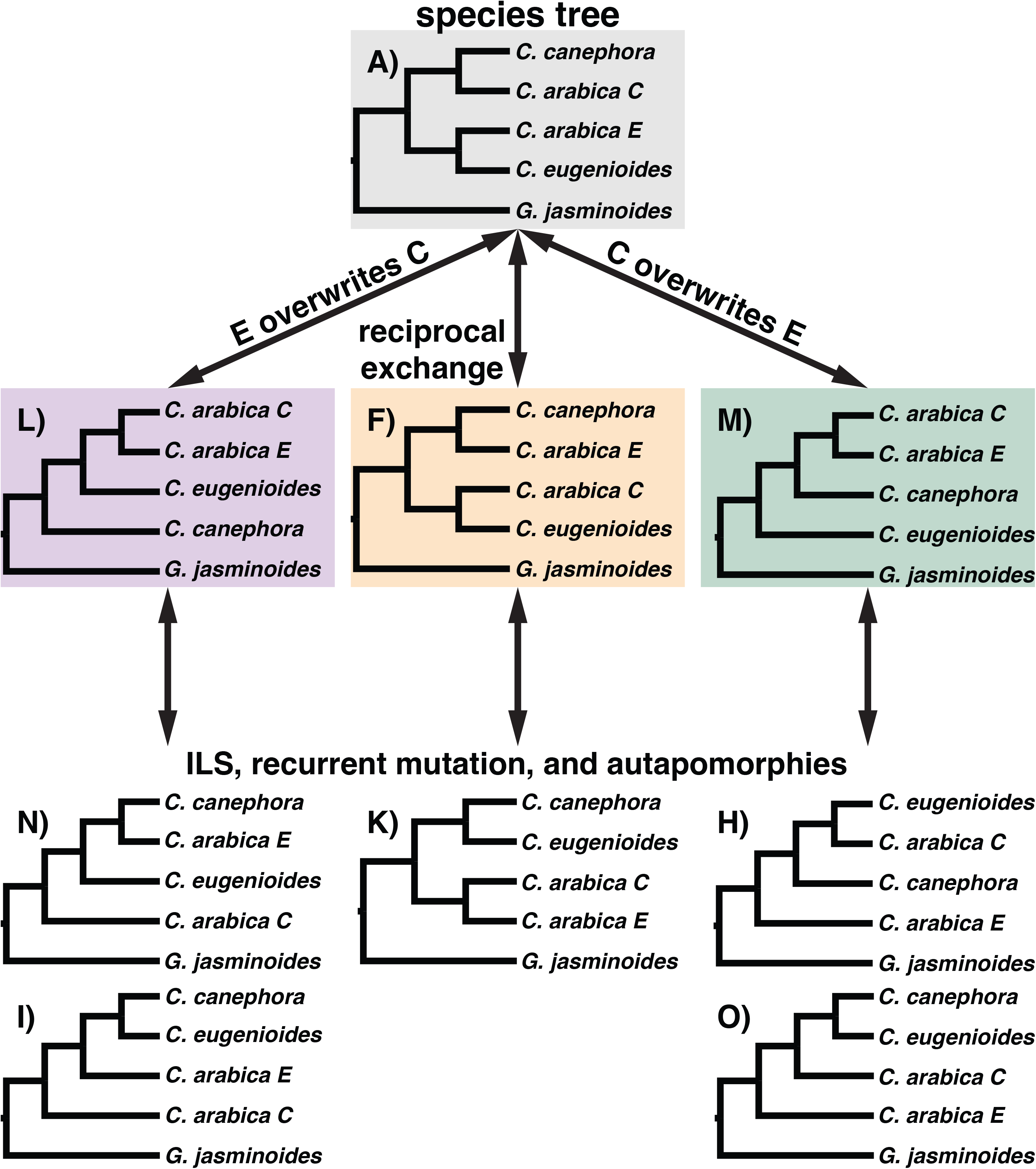
Comparing putative HGF gene trees with alternative gene tree topologies that are due to random sorting of alleles, recurrent mutations, and autapomorphies.

**Figure S4.**
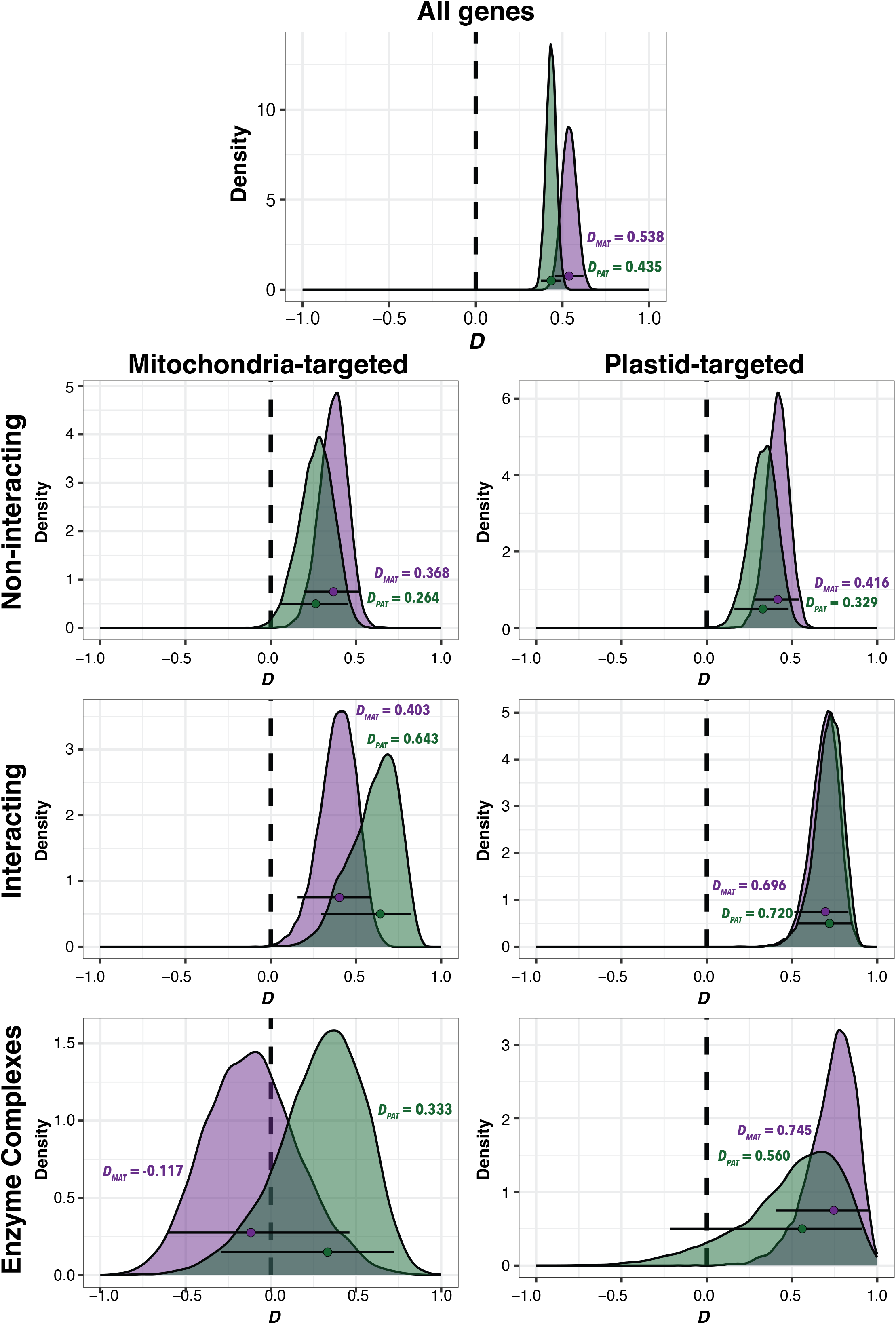
Reciprocal *D*-statistic estimates from Gblocks-trimmed alignments of the E subgenome overwriting the C subgenome (purple, *D_MAT_*) and of the C subgenome overwriting the E subgenome (green, *D_PAT_*) all genes (top panel) and in genes targeted to the mitochondria (left) and to the plastid (right). Genes are grouped according to the intimacy of interaction: non-interacting – top panels, interacting – middle panels, enzyme complexes – bottom panels. Points represent overall *D*-statistics, density plots depict distributions from 10,000 gene-level bootstrap replicates, and error bars represent 95% CIs. Distributions that are significantly greater than 0 are indicative of HGF in that direction.

## REFERENCES

Ågren, J. A., H.-R. Huang, and S. I. Wright. 2016. Transposable element evolution in the allotetraploid *Capsella bursa-pastoris*. American journal of botany 103: 1197–1202.

Akagi, T., I. M. Henry, T. Kawai, L. Comai, and R. Tao. 2016. Epigenetic regulation of the sex determination gene MeGI in polyploid persimmon. The Plant cell 28: 2905–2915.

Akama, S., R. Shimizu-Inatsugi, K. K. Shimizu, and J. Sese. 2014. Genome-wide quantification of homeolog expression ratio revealed nonstochastic gene regulation in synthetic allopolyploid *Arabidopsis*. Nucleic acids research 42: e46.

Baduel, P., L. Quadrana, B. Hunter, K. Bomblies, and V. Colot. 2019. Relaxed purifying selection in autopolyploids drives transposable element over-accumulation which provides variants for local adaptation. Nature communications 10: 5818.

Beck, E. A., A. C. Thompson, J. Sharbrough, E. Brud, and A. Llopart. 2015. Gene flow between *Drosophila yakuba* and *Drosophila santomea* in subunit V of cytochrome c oxidase: A potential case of cytonuclear cointrogression. Evolution; international journal of organic evolution 69: 1973–1986.

Bertioli, D. J., J. Jenkins, J. Clevenger, O. Dudchenko, D. Gao, G. Seijo, S. C. M. Leal-Bertioli, et al. 2019. The genome sequence of segmental allotetraploid peanut *Arachis hypogaea*. Nature genetics 51: 877–884.

Bomblies, K. 2020. When everything changes at once: finding a new normal after genome duplication. Proceedings. Biological sciences / The Royal Society 287: 20202154.

Bomblies, K., G. Jones, C. Franklin, D. Zickler, and N. Kleckner. 2016. The challenge of evolving stable polyploidy: could an increase in ‘crossover interference distance’ play a central role? Chromosoma 125: 287–300.

Burton, R. S., and F. S. Barreto. 2012. A disproportionate role for mtDNA in Dobzhansky-Muller incompatibilities? Molecular ecology 21: 4942–4957.

Cai, X., L. Chang, T. Zhang, H. Chen, L. Zhang, R. Lin, J. Liang, et al. 2021. Impacts of allopolyploidization and structural variation on intraspecific diversification in *Brassica rapa*. Genome biology 22: 166.

Campuzano-Duque, L. F., J. C. Herrera, C. Ged, and M. W. Blair. 2021. Bases for the establishment of robusta coffee (*Coffea canephora*) as a new crop for Colombia. Agronomy 11: 2550.

Camus, M. F., B. Alexander-Lawrie, J. Sharbrough, and G. D. D. Hurst. 2022. Inheritance through the cytoplasm. Heredity 129: 31–43.

Castresana, J. 2000. Selection of conserved blocks from multiple alignments for their use in phylogenetic analysis. Molecular biology and evolution 17: 540–552.

Chalhoub, B., F. Denoeud, S. Liu, I. A. P. Parkin, H. Tang, X. Wang, J. Chiquet, et al. 2014. Plant genetics. Early allopolyploid evolution in the post-Neolithic *Brassica napus* oilseed genome. Science 345: 950–953.

Chen, J., E. Li, X. Zhang, X. Dong, L. Lei, W. Song, H. Zhao, and J. Lai. 2017. Genome-wide nucleosome occupancy and organization modulates the plasticity of gene transcriptional status in maize. Molecular plant 10: 962–974.

Chen, S., F. Ren, L. Zhang, Y. Liu, X. Chen, Y. Li, L. Zhang, et al. 2018. Unstable allotetraploid tobacco genome due to frequent homeologous recombination, segmental deletion, and chromosome loss. Molecular plant 11: 914–927.

Chester, M., J. P. Gallagher, V. V. Symonds, A. V. Cruz da Silva, E. V. Mavrodiev, A. R. Leitch, P. S. Soltis, and D. E. Soltis. 2012. Extensive chromosomal variation in a recently formed natural allopolyploid species, *Tragopogon miscellus* (Asteraceae). Proceedings of the National Academy of Sciences of the United States of America 109: 1176–1181.

Chu, Y., D. Bertioli, C. M. Levinson, H. T. Stalker, C. C. Holbrook, and P. Ozias-Akins. 2021. Homoeologous recombination is recurrent in the nascent synthetic allotetraploid *Arachis ipaënsis × Arachis correntina* 4x and its derivatives. G3 11.

Comai, L. 2005. The advantages and disadvantages of being polyploid. Nature reviews. Genetics 6: 836–846.

Combes, M.-C., A. Dereeper, D. Severac, B. Bertrand, and P. Lashermes. 2013. Contribution of subgenomes to the transcriptome and their intertwined regulation in the allopolyploid *Coffea arabica* grown at contrasted temperatures. The New phytologist 200: 251–260.

Conover, J. L., and J. F. Wendel. 2022. Deleterious mutations accumulate faster in allopolyploid than diploid cotton (*Gossypium*) and unequally between subgenomes. Molecular biology and evolution 39.

Cros, J., M. C. Combes, P. Trouslot, F. Anthony, S. Hamon, A. Charrier, and P. Lashermes. 1998. Phylogenetic analysis of chloroplast DNA variation in *Coffea* L. Molecular phylogenetics and evolution 9: 109–117.

Deb, S. K., P. P. Edger, J. C. Pires, and M. R. McKain. 2023. Patterns, mechanisms, and consequences of homoeologous exchange in allopolyploid angiosperms: a genomic and epigenomic perspective. The New phytologist 238: 2284–2304.

Ding, M., and Z. J. Chen. 2018. Epigenetic perspectives on the evolution and domestication of polyploid plant and crops. Current opinion in plant biology 42: 37–48.

Douglas, G. M., G. Gos, K. A. Steige, A. Salcedo, K. Holm, E. B. Josephs, R. Arunkumar, et al. 2015. Hybrid origins and the earliest stages of diploidization in the highly successful recent polyploid *Capsella bursa-pastoris*. Proceedings of the National Academy of Sciences of the United States of America 112: 2806–2811.

Doyle, J. J., and J. E. Coate. 2019. Polyploidy, the nucleotype, and novelty: the impact of genome doubling on the biology of the cell. International journal of plant sciences.

Doyle, J. J., L. E. Flagel, A. H. Paterson, R. A. Rapp, D. E. Soltis, P. S. Soltis, and J. F. Wendel. 2008. Evolutionary genetics of genome merger and doubling in plants. Annual review of genetics 42: 443–461.

Drummond, D. A., J. D. Bloom, C. Adami, C. O. Wilke, and F. H. Arnold. 2005. Why highly expressed proteins evolve slowly. Proceedings of the National Academy of Sciences of the United States of America 102: 14338–14343.

Durand, E. Y., N. Patterson, D. Reich, and M. Slatkin. 2011. Testing for ancient admixture between closely related populations. Molecular biology and evolution 28: 2239–2252.

Edger, P. P., R. Smith, M. R. McKain, and A. M. Cooley. 2017. Subgenome dominance in an interspecific hybrid, synthetic allopolyploid, and a 140-year-old naturally established neo-allopolyploid monkeyflower. The Plant.

Edwards, K. D., N. Fernandez-Pozo, K. Drake-Stowe, M. Humphry, A. D. Evans, A. Bombarely, F. Allen, et al. 2017. A reference genome for *Nicotiana tabacum* enables map-based cloning of homeologous loci implicated in nitrogen utilization efficiency. BMC genomics 18: 448.

Ellis, R. J. 1979. The most abundant protein in the world. Trends in biochemical sciences 4: 241–244.

Endrizzi, J. E. 1962. The diploid-like cytological behavior of tetraploid cotton. Evolution; international journal of organic evolution 16: 325–329.

Fernandes Gyorfy, M. E. R. Miller, J. L. Conover, C. E. Grover, J. F. Wendel, D. B. Sloan, and J. Sharbrough. 2021. Nuclear-cytoplasmic balance: whole genome duplications induce elevated organellar genome copy number. The Plant journal: for cell and molecular biology.

Forsythe, E. S., C. E. Grover, E. R. Miller, J. L. Conover, M. A. Arick 2nd, M. C. F. Chavarro, S. C. M. Leal-Bertioli, et al. 2022. Organellar transcripts dominate the cellular mRNA pool across plants of varying ploidy levels. Proceedings of the National Academy of Sciences of the United States of America 119: e2204187119.

Forsythe, E. S., A. D. L. Nelson, and M. A. Beilstein. 2020. Biased gene retention in the face of introgression obscures species relationships. Genome biology and evolution 12: 1646–1663.

Forsythe, E. S., J. Sharbrough, J. C. Havird, J. M. Warren, and D. B. Sloan. 2019. CyMIRA: The cytonuclear molecular interactions reference for *Arabidopsis*. Genome biology and evolution 11: 2194–2202.

Fox, D. T., D. E. Soltis, P. S. Soltis, T.-L. Ashman, and Y. Van de Peer. 2020. Polyploidy: A biological force from cells to ecosystems. Trends in cell biology 30: 688–694.

Fulneček, J., R. Matyášek, and A. Kovařík. 2009. Faithful inheritance of cytosine methylation patterns in repeated sequences of the allotetraploid tobacco correlates with the expression of DNA methyltransferase gene families from both parental genomes. Molecular genetics and genomics: MGG 281: 407–420.

Gaeta, R. T., J. C. Pires, F. Iniguez-Luy, E. Leon, and T. C. Osborn. 2007. Genomic changes in resynthesized *Brassica napus* and their effect on gene expression and phenotype. The Plant cell 19: 3403–3417.

Gong, L., M. Olson, and J. F. Wendel. 2014. Cytonuclear evolution of rubisco in four allopolyploid lineages. Molecular biology and evolution 31: 2624–2636.

Gong, L., A. Salmon, M.-J. Yoo, K. K. Grupp, Z. Wang, A. H. Paterson, and J. F. Wendel. 2012. The cytonuclear dimension of allopolyploid evolution: an example from cotton using rubisco. Molecular biology and evolution 29: 3023–3036.

Gonzalo, A., P. Parra-Nunez, A. L. Bachmann, E. Sanchez-Moran, and K. Bomblies. 2023. Partial cytological diploidization of neoautotetraploid meiosis by induced cross-over rate reduction. Proceedings of the National Academy of Sciences of the United States of America 120: e2305002120.

Gordon, S. P., B. Contreras-Moreira, J. J. Levy, A. Djamei, A. Czedik-Eysenberg, V. S. Tartaglio, A. Session, et al. 2020. Gradual polyploid genome evolution revealed by pan-genomic analysis of *Brachypodium hybridum* and its diploid progenitors. Nature communications 11: 3670.

Guo, H., X. Wang, H. Gundlach, K. F. X. Mayer, D. G. Peterson, B. E. Scheffler, P. W. Chee, and A. H. Paterson. 2014. Extensive and biased intergenomic nonreciprocal DNA exchanges shaped a nascent polyploid genome, *Gossypium* (cotton). Genetics 197: 1153– 1163.

Havird, J. C., N. S. Whitehill, C. D. Snow, and D. B. Sloan. 2015. Conservative and compensatory evolution in oxidative phosphorylation complexes of angiosperms with highly divergent rates of mitochondrial genome evolution. Evolution; international journal of organic evolution 69: 3069–3081.

Heslop-Harrison, J. S. P., T. Schwarzacher, and Q. Liu. 2023. Polyploidy: its consequences and enabling role in plant diversification and evolution. Annals of botany 131: 1–10.

Hibbins, M. S., and M. W. Hahn. 2022. Corrigendum to: phylogenomic approaches to detecting and characterizing introgression. Genetics 220.

Holm, S. 1979. A Simple Sequentially Rejective multiple test procedure. Scandinavian journal of statistics, theory and applications 6: 65–70.

Hu, G., R. Hovav, C. E. Grover, A. Faigenboim-Doron, N. Kadmon, J. T. Page, J. A. Udall, and J. F. Wendel. 2016. Evolutionary conservation and divergence of gene coexpression networks in *Gossypium* (cotton) seeds. Genome biology and evolution 8: 3765–3783.

Jarvis, D. E., Y. S. Ho, D. J. Lightfoot, S. M. Schmöckel, B. Li, T. J. A. Borm, H. Ohyanagi, et al. 2017. The genome of *Chenopodium quinoa*. Nature 542: 307–312.

Jiao, Y., N. J. Wickett, S. Ayyampalayam, A. S. Chanderbali, L. Landherr, P. E. Ralph, L. P. Tomsho, et al. 2011. Ancestral polyploidy in seed plants and angiosperms. Nature 473: 97– 100.

Katoh, K., and D. M. Standley. 2013. MAFFT multiple sequence alignment software version 7: improvements in performance and usability. Molecular biology and evolution 30: 772–780.

Kihara, H., and T. Ono. 1926. Chromosomenzahlen und systematische Gruppierung der Rumex-Arten. Zeitschrift für Zellforschung und Mikroskopische Anatomie 4: 475–481.

Kremnev, D., and A. Strand. 2014. Plastid encoded RNA polymerase activity and expression of photosynthesis genes required for embryo and seed development in *Arabidopsis*. Frontiers in plant science 5: 385.

Kühn, K., T. Obata, K. Feher, R. Bock, A. R. Fernie, and E. H. Meyer. 2015. Complete mitochondrial complex I deficiency induces an up-regulation of respiratory fluxes that is abolished by traces of functional complex I. Plant physiology 168: 1537–1549.

Landis, J. B., A. Kurti, A. J. Lawhorn, A. Litt, and E. W. McCarthy. 2020. Differential gene expression with an emphasis on floral organ size differences in natural and synthetic polyploids of *Nicotiana tabacum* (Solanaceae). Genes 11.

Lashermes, P., M. C. Combes, J. Robert, P. Trouslot, A. D’Hont, F. Anthony, and A. Charrier. 1999. Molecular characterisation and origin of the *Coffea arabica* L. genome. Molecular & general genetics: MGG 261: 259–266.

Lashermes, P., Y. Hueber, M.-C. Combes, D. Severac, and A. Dereeper. 2016. Inter-genomic DNA exchanges and homeologous gene silencing shaped the nascent allopolyploid coffee genome (*Coffea arabica* L.). G3 6: 2937–2948.

Leal-Bertioli, S. C. M., I. J. Godoy, J. F. Santos, J. J. Doyle, P. M. Guimarães, B. L. Abernathy, S. A. Jackson, et al. 2018. Segmental allopolyploidy in action: Increasing diversity through polyploid hybridization and homoeologous recombination. American journal of botany 105: 1053–1066.

Leitch, A. R., and I. J. Leitch. 2008. Genomic plasticity and the diversity of polyploid plants. Science 320: 481–483.

Li, C., X. Wang, Y. Xiao, X. Sun, J. Wang, X. Yang, Y. Sun, et al. 2020. Co-evolution in hybrid genomes: nuclear-encoded rubisco small subunits and their plastid-targeting translocons accompanying sequential allopolyploidy events in *Triticum*. Molecular biology and evolution.

Madlung, A., R. W. Masuelli, B. Watson, S. H. Reynolds, J. Davison, and L. Comai. 2002. Remodeling of DNA methylation and phenotypic and transcriptional changes in synthetic *Arabidopsis* allotetraploids. Plant physiology 129: 733–746.

Mason, A. S., and J. C. Pires. 2015. Unreduced gametes: meiotic mishap or evolutionary mechanism? Trends in genetics: TIG 31: 5–10.

Mason, A. S., and J. F. Wendel. 2020. Homoeologous exchanges, segmental allopolyploidy, and polyploid genome evolution. Frontiers in genetics 11: 1014.

Oberprieler, C., M. Talianova, and J. Griesenbeck. 2019. Effects of polyploidy on the coordination of gene expression between organellar and nuclear genomes in *Leucanthemum* Mill. (Compositae, Anthemideae). Ecology and evolution 9: 9100–9110.

One Thousand Plant Transcriptomes Initiative. 2019. One thousand plant transcriptomes and the phylogenomics of green plants. Nature 574: 679–685.

Orantes-Bonilla, M., M. Makhoul, H. Lee, H. S. Chawla, P. Vollrath, A. Langstroff, F. J. Sedlazeck, et al. 2022. Frequent spontaneous structural rearrangements promote rapid genome diversification in a *Brassica napus* F1 generation. Frontiers in plant science 13: 1057953.

Oruganti, V., H. Toegelová, A. Pečinka, A. Madlung, and K. Schneeberger. 2023. Rapid large-scale genomic introgression in *Arabidopsis suecica* via an autoallohexaploid bridge. Genetics 223.

Otto, S. P. 2007. The evolutionary consequences of polyploidy. Cell 131: 452–462.

Otto, S. P., and J. Whitton. 2000. Polyploid incidence and evolution. Annual review of genetics.

Pont, C., T. Leroy, M. Seidel, A. Tondelli, W. Duchemin, D. Armisen, D. Lang, et al. 2019. Tracing the ancestry of modern bread wheats. Nature genetics 51: 905–911.

Ramírez-González, R. H., P. Borrill, D. Lang, S. A. Harrington, J. Brinton, L. Venturini, M. Davey, et al. 2018. The transcriptional landscape of polyploid wheat. Science 361.

Ramsey, J., and D. W. Schemske. 2003. Neopolyploidy in flowering plants.

Rand, D. M., R. A. Haney, and A. J. Fry. 2004. Cytonuclear coevolution: the genomics of cooperation. Trends in ecology & evolution 19: 645–653.

Rao, X., J. Ren, W. Wang, R. Chen, Q. Xie, Y. Xu, D. Li, et al. 2023. Comparative DNA-methylome and transcriptome analysis reveals heterosis- and polyploidy-associated epigenetic changes in rice. The Crop Journal 11: 427–437.

Raven, J. A. 2013. Rubisco: still the most abundant protein of Earth? The New phytologist 198: 1–3.

Rockenbach, K., J. C. Havird, J. G. Monroe, D. A. Triant, D. R. Taylor, and D. B. Sloan. 2016. Positive selection in rapidly evolving plastid-nuclear enzyme complexes. Genetics 204: 1507–1522.

Román-Palacios, C., C. A. Medina, S. H. Zhan, and M. S. Barker. 2021. Animal chromosome counts reveal a similar range of chromosome numbers but with less polyploidy in animals compared to flowering plants. Journal of evolutionary biology 34: 1333–1339.

Ruprecht, C., R. Lohaus, K. Vanneste, M. Mutwil, Z. Nikoloski, Y. Van de Peer, and S. Persson. 2017. Revisiting ancestral polyploidy in plants. Science advances 3: e1603195.

Salmon, A., M. L. Ainouche, and J. F. Wendel. 2005. Genetic and epigenetic consequences of recent hybridization and polyploidy in *Spartina* (Poaceae). Molecular ecology 14: 1163– 1175.

Salmon, A., L. Flagel, B. Ying, J. A. Udall, and J. F. Wendel. 2010. Homoeologous nonreciprocal recombination in polyploid cotton. The New phytologist 186: 123–134.

Salojarvi, J., A. Rambani, Z. Yu, R. Guyot, S. Strickler, M. Lepelley, C. Wang, et al. 2023. The genome and population genomics of allopolyploid *Coffea arabica* reveal the diversification history of modern coffee cultivars. bioRxiv: 2023.09.06.556570.

Scalabrin, S., L. Toniutti, G. Di Gaspero, D. Scaglione, G. Magris, M. Vidotto, S. Pinosio, et al. 2020. A single polyploidization event at the origin of the tetraploid genome of *Coffea arabica* is responsible for the extremely low genetic variation in wild and cultivated germplasm. Scientific reports 10: 4642.

Schiessl, S.-V., E. Katche, E. Ihien, H. S. Chawla, and A. S. Mason. 2019. The role of genomic structural variation in the genetic improvement of polyploid crops. The Crop Journal 7: 127–140.

Schnable, J. C., B. S. Pedersen, S. Subramaniam, and M. Freeling. 2011. Dose–sensitivity, conserved non-coding sequences, and duplicate gene retention through multiple tetraploidies in the grasses. Frontiers in plant science 2.

Sharbrough, J., J. L. Conover, M. Fernandes Gyorfy, C. E. Grover, E. R. Miller, J. F. Wendel, and D. B. Sloan. 2022. Global patterns of subgenome evolution in organelle-targeted genes of six allotetraploid angiosperms. Molecular biology and evolution 39.

Sharbrough, J., J. L. Conover, J. A. Tate, J. F. Wendel, and D. B. Sloan. 2017. Cytonuclear responses to genome doubling. American journal of botany 104: 1277–1280.

Shcherban, A. B., E. D. Badaeva, A. V. Amosova, I. G. Adonina, and E. A. Salina. 2008. Genetic and epigenetic changes of rDNA in a synthetic allotetraploid, Aegilops sharonensis x Ae. umbellulata. Genome / National Research Council Canada = Genome / Conseil national de recherches Canada 51: 261–271.

Sloan, D. B., J. C. Havird, and J. Sharbrough. 2017. The on-again, off-again relationship between mitochondrial genomes and species boundaries. Molecular ecology.

Sloan, D. B., J. M. Warren, A. M. Williams, Z. Wu, S. E. Abdel-Ghany, A. J. Chicco, and J. C. Havird. 2018. Cytonuclear integration and co-evolution. Nature reviews. Genetics 19: 635– 648.

Soltis, D. E., C. J. Visger, and P. S. Soltis. 2014. The polyploidy revolution then… and now: Stebbins revisited. American journal of botany 101: 1057–1078.

Song, M. J., B. I. Potter, J. J. Doyle, and J. E. Coate. 2020. Gene balance predicts transcriptional responses immediately following ploidy change in *Arabidopsis thaliana*. The Plant cell 32: 1434–1448.

Song, Q., T. Zhang, D. M. Stelly, and Z. J. Chen. 2017. Epigenomic and functional analyses reveal roles of epialleles in the loss of photoperiod sensitivity during domestication of allotetraploid cottons. Genome biology 18: 99.

Stamatakis, A. 2014. RAxML version 8: a tool for phylogenetic analysis and post-analysis of large phylogenies. Bioinformatics 30: 1312–1313.

Steenwyk, J. L., T. J. Buida 3rd, Y. Li, X.-X. Shen, and A. Rokas. 2020. ClipKIT: A multiple sequence alignment trimming software for accurate phylogenomic inference. PLoS biology 18: e3001007.

Stein, A., O. Coriton, M. Rousseau-Gueutin, B. Samans, S. V. Schiessl, C. Obermeier, I. A. P. Parkin, et al. 2017. Mapping of homoeologous chromosome exchanges influencing quantitative trait variation in *Brassica napus*. Plant biotechnology journal 15: 1478–1489.

Team, R. C., and R. Core Team. R: a language and environment for statistical computing. R Found. Stat Comput Vienna Austria.

Wendel, J. F. 2015. The wondrous cycles of polyploidy in plants. American journal of botany 102: 1753–1756.

Wickham, H. 2011. ggplot2. Wiley interdisciplinary reviews. Computational statistics 3: 180– 185.

Williams, A. M., G. Friso, K. J. van Wijk, and D. B. Sloan. 2019. Extreme variation in rates of evolution in the plastid Clp protease complex. The Plant journal: for cell and molecular biology 98: 243–259.

Wolfe, K. H., W. H. Li, and P. M. Sharp. 1987. Rates of nucleotide substitution vary greatly among plant mitochondrial, chloroplast, and nuclear DNAs. Proceedings of the National Academy of Sciences of the United States of America 84: 9054–9058.

Xiong, Z., R. T. Gaeta, P. P. Edger, Y. Cao, K. Zhao, S. Zhang, and J. C. Pires. 2020. Chromosome inheritance and meiotic stability in allopolyploid *Brassica napus*. G3 Genes|Genomes|Genetics.

Yang, J., D. Liu, X. Wang, C. Ji, F. Cheng, B. Liu, Z. Hu, et al. 2016. The genome sequence of allopolyploid *Brassica juncea* and analysis of differential homoeolog gene expression influencing selection. Nature genetics 48: 1225–1232.

Yang, J.-R., B.-Y. Liao, S.-M. Zhuang, and J. Zhang. 2012. Protein misinteraction avoidance causes highly expressed proteins to evolve slowly. Proceedings of the National Academy of Sciences of the United States of America 109: E831–40.

Yoo, M.-J., E. Szadkowski, and J. F. Wendel. 2013. Homoeolog expression bias and expression level dominance in allopolyploid cotton. Heredity 110: 171–180.

Yuan, Y., Z. Milec, P. E. Bayer, J. Vrána, J. Doležel, D. Edwards, W. Erskine, and P. Kaur. 2018. Large-scale structural variation detection in subterranean clover subtypes using optical mapping. Frontiers in plant science 9: 971.

Zhang, Z., X. Gou, H. Xun, Y. Bian, X. Ma, J. Li, N. Li, et al. 2020. Homoeologous exchanges occur through intragenic recombination generating novel transcripts and proteins in wheat and other polyploids. Proceedings of the National Academy of Sciences of the United States of America 117: 14561–14571.

